# The pathological hallmarks of Alzheimer’s disease derive from compensatory responses to NMDA receptor insufficiency

**DOI:** 10.1101/418566

**Authors:** Selina Sohre, Bernd Moosmann

## Abstract

Alzheimer’s disease is characterized by intracellular aggregates of hyperphosphorylated tau protein and extracellular plaques of amyloid β peptide, a product of APP processing. The origin of these pathological hallmarks has remained elusive. Here, we have tested the idea that both alterations, at the onset of the disease, may constitute compensatory responses to the same causative and initial trigger, namely NMDA receptor insufficiency. Treatment of rat cortical neurons with the specific NMDA receptor antagonist AP5 within 4 h caused a significant increase in tau phosphorylation at the AT8 and S404 epitopes as well as an increase in APP expression and Aβ _40_ secretion. Single intraperitoneal injections of the NMDA receptor open channel blocker MK-801 into wild-type mice reproduced all of these changes in a brain region-specific fashion either at latency 4 h or 24 h. Subchronic treatment with MK-801 for 6 weeks induced AT8, S404 and S396 immunoreactivity selectively in female mice. We conclude that the pivotal pathological alterations in Alzheimer’s disease represent runaway physiological responses to persistently insufficient excitatory neurotransmission. In view of the evidence for excitatory insufficiency in trisomy 21 patients, PS1 mutation carriers and ApoE4 carriers, our data suggest a common pathomechanism behind familial, sporadic, and risk allele-triggered Alzheimer’s disease. The potential of this mechanism to reconcile previous conflicting observations is discussed.

## Introduction

Despite intense efforts, Alzheimer’s disease (AD) has remained an insufficiently understood disorder, which is evidenced by an extensive series of failures of drug candidates in clinical trials (Braak et al., 1999; Cummings et al., 2014; Herrup, 2015).

In general pathology, microscopically visible hallmarks of a disease often reflect exaggerated physiological responses to an invisible disease trigger rather than representing the disease origin by themselves. Frequently, these physiological responses are compensatory in nature, i.e. directed at the counterbalancing of the effect evoked by the original disease trigger or its removal. For example, in a typical form of serum amyloid A (SAA) amyloidosis, chronic infection with tuberculosis leads to a massive, but fruitless compensatory secretion of the anti-infectious SAA protein by the liver, which precedes and determines its erratic and often fatal deposition as cleaved amyloid A protein in kidney and gut (Westermark et al., 2014).

Translating this reasoning to the hallmarks of AD, the idea was pursued that both APP expression and amyloid β (Aβ) formation as well as tau phosphorylation may constitute initially purposeful physiological responses to a common, but unknown molecular trigger. If that trigger persisted nevertheless, both responses would become more and more enforced, with the result that exaggerated and eventually toxic concentrations were attained in the end.

In distilling the literature for the most likely such trigger as based on today’s knowledge, we reached the conclusion that chronic antagonism of excitatory neurotransmission via NMDA receptors would be the most plausible candidate. In brief, this conclusion was drawn on the basis of the following evidence.

### Amyloid

The most thoroughly described property of Aβ peptides is their induction of excitotoxicity at supraphysiological doses ranging from 10^-7^ M to 10^-4^ M, mostly after pharmacological application in vitro or mutant APP overexpression in vivo (Mattson et al., 1992; Harkany et al., 2000; Danysz and Parsons, 2012). Supposing that this potent excitotoxic activity at supraphysiological doses was a mirror image of Aβ’s physiological role at 10^-10^ M to 10^-9^ M concentrations, the “maximum parsimony” hypothesis was derived that physiological Aβ should be viewed as glutamatergic sensitizer, probably mediated through NMDA receptors. In fact, work from three independent laboratories has shown that physiological, picomolar concentrations of Aβ rapidly enhance NMDA receptor-dependent long-term potentiation (LTP) in vitro and improve learning and memory formation in vivo (Puzzo et al., 2008; Garcia-Osta and Alberini, 2009; Morley et al., 2010). Supraphysiological, transgenic concentrations of Aβ have often been seen to later elicit the opposite effect, probably due to enhanced NMDA receptor internalization (Snyder et al., 2005) in an effort to prevent looming excitotoxicity.

### Tau

Tau binds to and stabilizes microtubules (Wang and Mandelkow, 2016), but also physically impairs kinesin migration (Dixit et al., 2008) and thus anterograde transport of newly synthesized APP (Stamer et al., 2002) and NMDA receptors (Setou et al., 2000). Phosphorylation generally dissociates tau from microtubules, potentially facilitating protein transport, but at the danger of microtubule disintegration (Dixit et al., 2008; Wang and Mandelkow, 2016). Moreover, tau phosphorylation has been reported to entail its redistribution into dendritic structures (Hoover et al., 2010), where it is thought to sensitize NMDA receptors through phosphorylation by Src kinase fyn (Ittner et al., 2010). Supraphysiological tau accumulation in dendritic spines might then induce excitotoxicity (Ittner et al., 2010) or enhanced receptor internalization (Hoover et al., 2010). Notably, tau reduction has been described to suppress seizure activity and hyperexcitability in different disease models in vivo (Roberson et al., 2007; Holth et al., 2013) and to diminish glutamate excitotoxicity in vitro (Miyamoto et al., 2017). Hence, tau protein and its activated, redistributed phospho-forms appear to physiologically enhance excitatory neurotransmission, which implies that glutamatergic insufficiency might induce tau phosphorylation as a normal homeostatic response.

Therefore, the effect of a selective pharmacological blockade of NMDA receptors was tested under fully physiological, non-transgenic conditions in vitro and in vivo. It was found that this intervention led to a rapid and concomitant induction of APP expression, Aβ formation and tau phosphorylation on AD-related epitopes.

## Results

Cortical neurons from E17 Sprague-Dawley rats were prepared, cultivated to maturity in Neurobasal medium for 14-17 days and then transferred to defined physiological Ringer solution for experimental investigation. Spiking activity under comparable conditions (Sun et al., 2010) arose at around day 7 and started to plateau at day 12, whereas coordinated bursting activity reached a stable maximum after day 10 (Sun et al., 2010). Consequently, investigations dependent on spontaneous neuronal activity were deemed feasible in these cultures. Since depolarization with 25 mM potassium (K^+^) is known to reversibly increase spontaneous NMDA receptor-dependent activity in vitro (Heck et al., 2008), all experiments were done under standard (2.5 mM K^+^) as well as depolarizing (27.5 mM K^+^) conditions.

Treatment of these cortical cultures with 100 µM AP5, a highly specific competitive NMDA receptor antagonist (Watkins et al., 1990), within 4 h of incubation induced a significant increase in tau phosphorylation at the AT8, S404 and S396 epitopes (Fig. 1a-c). Separate incubation with glutamatergic agonists, either 5 µM NMDA or 5 µM L-glutamate, did not significantly reduce tau phosphorylation below the baseline level, however (Fig. 1b,c). Consistent with the NMDA receptor binding affinities of the employed compounds, the effect of co-incubation with these agonists and AP5 was dominated by the presence of AP5 (Fig. 1b,c). K^+^-induced depolarization had no visible influence on the degree of tau phosphorylation, potentially due to the absence of the NMDA receptor blocker Mg^2+^ in the measurement Ringer solution. Combined analysis of six independent cell cultures, three for each K^+^ concentration, yielded a significance level of p2 < 0.001 (by Two-way ANOVA) for the induction of tau phosphorylation by AP5 at all three investigated phospho-sites (Fig. 1c). Our data concur with earlier reports of an NMDA receptor mediated regulation of tau phosphorylation in rat cortical (Fleming and Johnson, 1995) and hippocampal (Allyson et al., 2010; De Montigny et al., 2013) slices.

**Figure 1.**
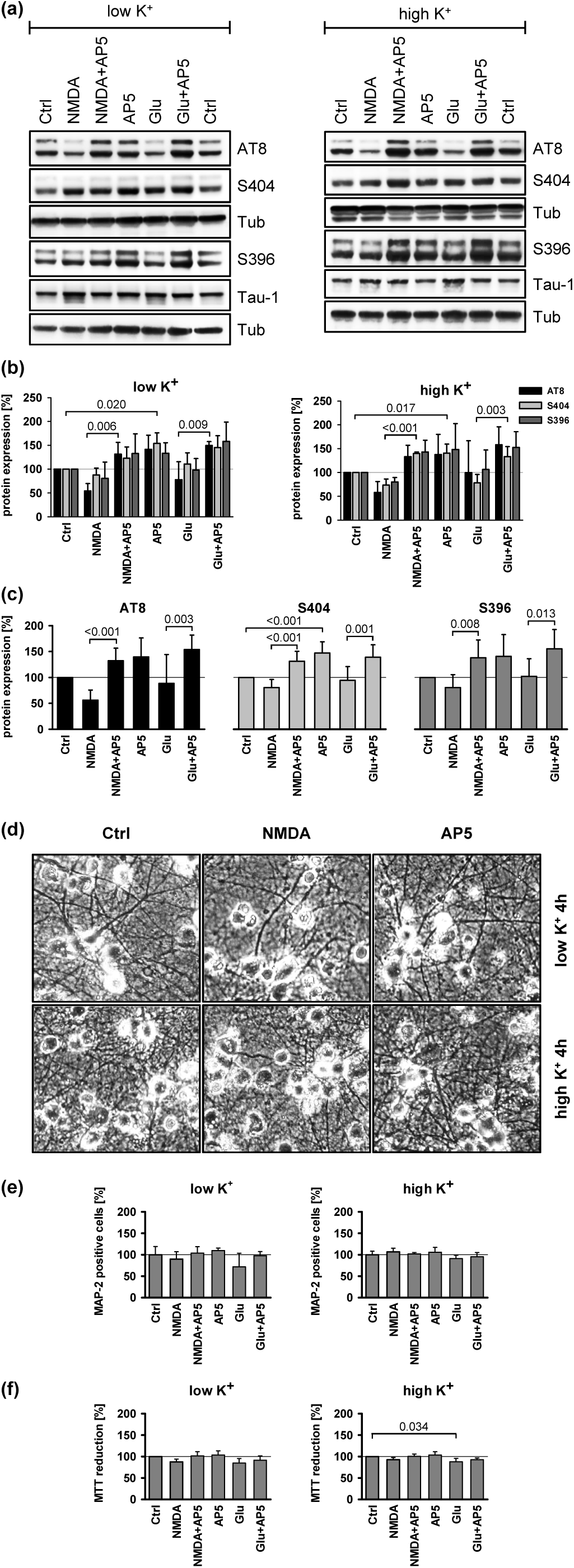
Tau phosphorylation in rat cortical neurons. **(a)** Effect of NMDA (5 µM), glutamate (Glu, 5 µM) and AP5 (100 µM) treatment on epitope-specific tau phosphorylation at the AT8, S404 and S396 sites in rat cortical neurons maintained under low (2.5 mM) and high (27.5 mM) K^+^ conditions for 4 h. The Tau-1 antibody recognizes unphosphorylated total tau; tubulin (Tub) was used as loading control. Tub control blots always refer to the 1-2 blots directly above. **(b)** Quantification of n = 3 independent Western blot experiments performed as in (a). After individual normalization to Tub, signal intensities of each blot were related to accordingly normalized total tau. Statistical evaluations were done by Two-way ANOVA with factor #1 being agonist exposure (NMDA or Glu) and factor #2 being antagonist (AP5) exposure. Here and throughout this work, displayed significance levels result from pairwise post-hoc tests executed only in case of ANOVA significance (px < 0.05); further details are provided in the Supplemental information. ANOVA significance levels were: Anti-AT8 (low K+: p_1_= 0.246, p_2_ < 0.001, n = 3; high K+: p_1_ = 0.473, p_2_ = 0.024, n = 3), anti-S404 (low K+: p_1_ = 0.241, p_2_ = 0.004, n = 3; high K+: p_1_ = 0.322, p_2_ < 0.001, n = 3) and anti-S396 (low K+: p_1_ = 0.625, p_2_ = 0.017, n = 3; high K+: p_1_ = 0.735, p_2_ = 0.017, n = 3). **(c)** Combined statistical analysis of tau phosphorylation under low and high K^+^ conditions. Anti-AT8: p_1_ = 0.122, p_2_ < 0.001, n = 6; anti-S404: p_1_ = 0.148, p_2_ < 0.001, n = 6; anti-S396: p_1_ = 0.399, p_2_ < 0.001, n = 6. **(d)** Phase-contrast images of cortical neurons treated with 5 µM NMDA or 100 µM AP5 under low and high K^+^ conditions for 4 h as in the above experiments. **(e)** Assessment of neuronal survival by counting MAP-2 positive cells. Cultures treated as in (a)-(d) were stained for neuron-specific MAP-2 immunoreactivity, photographed and examined quantitatively. For each biological replicate (n), neuron counts from three different pictures were added and normalized to the total number of plated cells in the same areas as per DAPI staining. Low K^+^: p_1_ = 0.110, p_2_ = 0.039, n = 4; high K+: p_1_ = 0.075, p_2_ = 0.659, n = 4. **(f)** Assessment of metabolic activity in corresponding cultures by MTT reduction. Each biological replicate (n) was derived from 5 replicates on the same multi-well plate. Low K+: p_1_ = 0.042, p_2_ = 0.067, n = 4; high K+: p_1_ = 0.009, p_2_ = 0.063, n = 4.

To monitor the potential role of cytotoxicity and metabolic modulation by the employed glutamate receptor ligands, the cells were examined microscopically (Fig. 1d), immunostained for the neuron-specific marker protein MAP-2 (Fig. 1e), and they were analyzed by the MTT metabolic activity assay (Fig. 1f). Irrespective of the treatment group, cells were morphologically indistinguishable from the control after 4 h (Fig. 1d). Quantification of MAP-2 positive cells did not reveal any differences under high K+ conditions, albeit a significant effect (p2 < 0.039) towards higher cell numbers in AP5 treated groups was seen in low K^+^ cultures (Fig. 1e). Conversely, AP5 treatment had no significant effect in the MTT metabolic activity assay, whereas glutamatergic agonist treatment involved a significant reduction in MTT conversion (p1 = 0.042 for low K+, p1 = 0.009 for high K+) (Fig. 1f). As evident from Fig. 1e and 1f, modulatory effect sizes were general small and cannot explain the effects of AP5 in the above tau phosphorylation experiments. Still, the glutamate-alone cultures may have been influenced to some extent by early excitotoxicity or metabotropic glutamate receptor ligation, with unknown consequences as regards tau phosphorylation.

The above-described 4 h cortical cultures were furthermore analyzed for APP expression and Aβ secretion. Treatment with AP5 evoked a highly significant induction of APP expression (Fig. 2a,b) in low and high K+ cultures (p_2_ < 0.001 in both cases), whereas glutamatergic agonist treatment had no significant effect as per Two-way ANOVA. Nevertheless, a lower average expression of APP in the agonist-alone treated cultures was noted (Fig. 2a,b). A combined evaluation of APP expression in all six independent cell cultures is provided in Fig. 2c. Analysis of these six cultures for Aβ secretion demonstrated a significant induction of Aβ_40_ formation by AP5 treatment (p_2_ = 0.003), whereas Aβ_42_ levels were unaffected (Fig. 2d). Hence, inhibition of excitatory communication via NMDA receptors in vitro evokes a rapid and concomitant response of tau phosphorylation, APP expression and Aβ formation.

**Figure 2.**
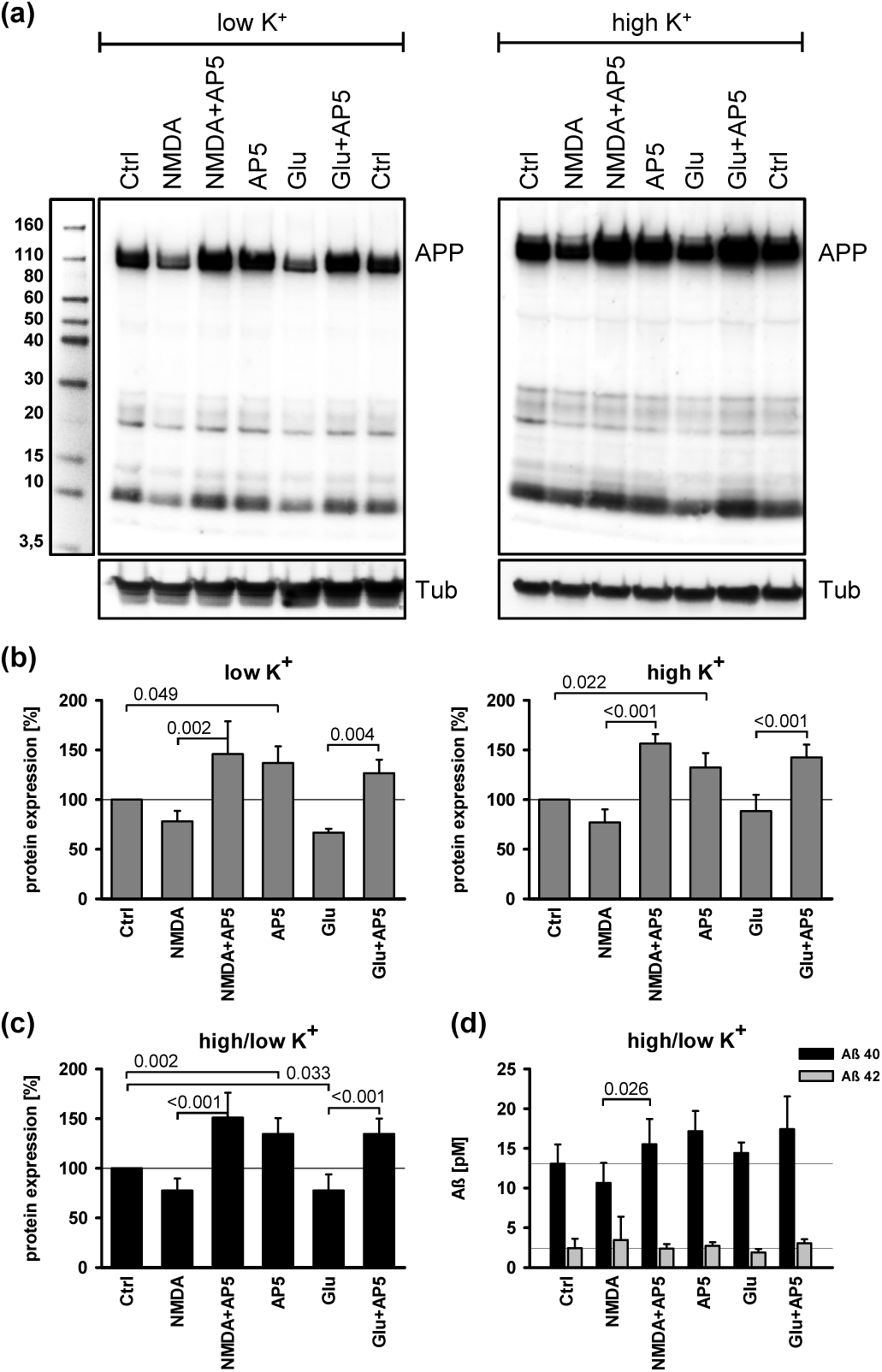
APP expression and processing in rat cortical neurons. **(a)** Effect of NMDA (5 µM), glutamate (Glu, 5 µM) and AP5 (100 µM) on APP expression in rat cortical neurons maintained under low (2.5 mM) and high (27.5 mM) K^+^ conditions for 4 h. The antibody recognizes the first 10 amino acids of the Aβ sequence. **(b)** Quantification of n = 3 independent Western blots performed as in (a) under low and high K^+^ conditions. Statistical evaluation was done by Two-way ANOVA as in Fig. 1b; significance levels indicated in the graph result from post-hoc tests. Low K+: p1 = 0.209, p2 < 0.001, n = 3; high K+: p_1_ = 0.988, p_2_ < 0.001, n = 3. **(c)** Combined analysis for both cultivation conditions: p_1_ = 0.270, p_2_ < 0.001, pinteraction = 0.034, n = 6. **(d)** Soluble Aβ_40_ and Aβ_42_ levels in the supernatant of cell cultures treated as above were analyzed by ELISA. Depicted is the combined analysis of the low and high K^+^ cultures. Aβ_40_: p_1_ = 0.100, p_2_ = 0.003, n = 6; Aβ_42_: p_1_ = 0.784, p_2_ = 0.925, n = 6.

Wild-type male “black 6” mice (C57BL/6JRj from Janvier) at an age of 10 weeks were chosen to investigate the behavior of these molecules after reversible NMDA receptor blockade in vivo. For this purpose, the BBB-permeable and specific NMDA antagonist MK-801 was injected intraperitoneally (ip) at a concentration of 1.0 mg/kg, followed by stereotactic brain tissue preparation after 4 h or 24 h. In accordingly treated rats, MK-801 in the brain peaked within 10-30 min after ip injection and was eliminated with a half-life of 2 h (Vezzani et al., 1989), which corresponds to the observed hyperactivity phenotype of the here studied mice that resolved after approximately 2 h. Analysis of tissue-specific tau phosphorylation (Fig. 3) indicated that the S404 epitope was significantly induced in the hippocampus (p = 0.022 by One-way ANOVA) and the cortex (p = 0.016).Moreover, AT8 phosphorylation was induced in the cortex after 24 h (p = 0.003). The thalamus did not exhibit any changes in response to MK-801 treatment (Fig. 3e).

**Figure 3.**
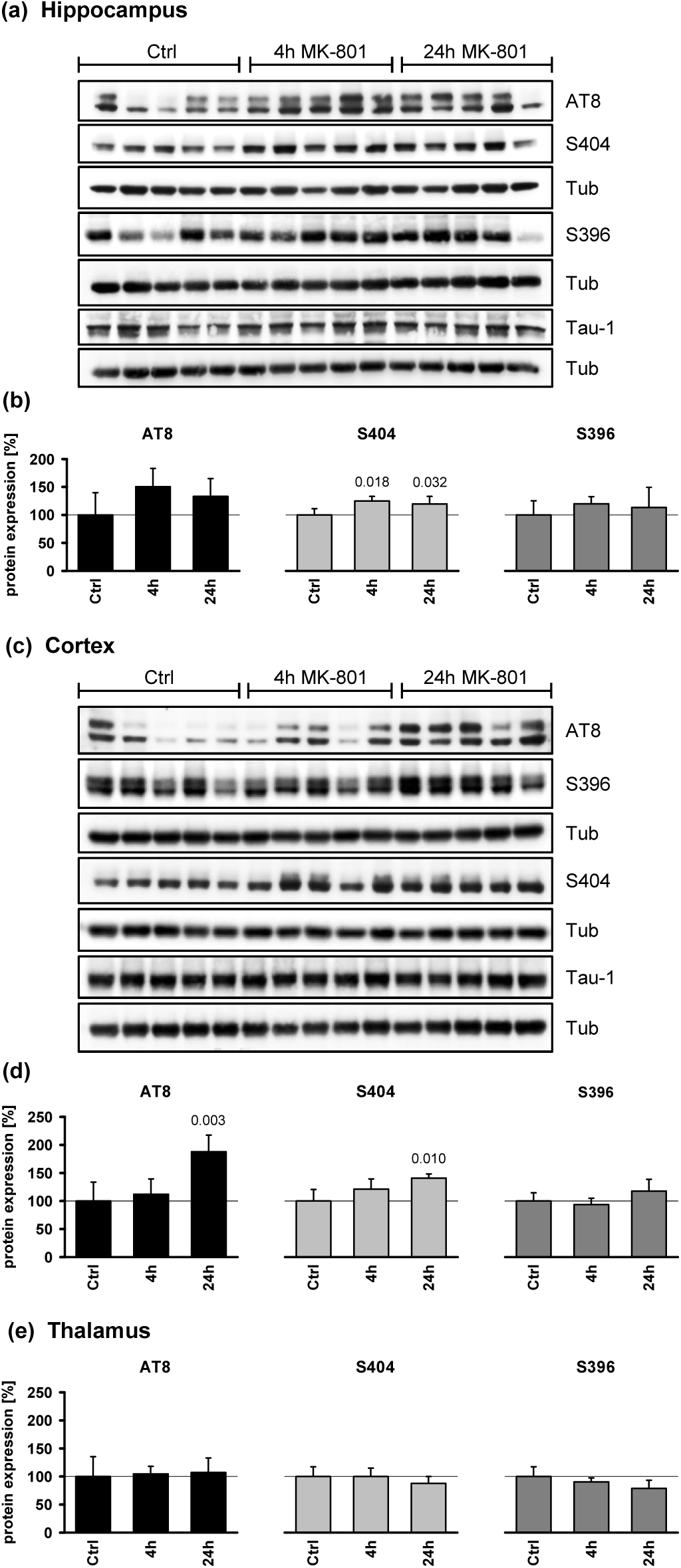
Tau phosphorylation in MK-801 treated mice. Male mice treated with MK-801 (1.0 mg/kg ip) were analyzed after 4 h and 24 h for specific tau phosphorylation. The control group was exposed to saline for 4 h. Different brain regions were stereotactically dissected as described in the Methods. Expression levels were normalized blot-wise to Tub and afterwards related to total tau expression as in Fig. 1. The three groups (n = 5) were evaluated by One-way ANOVA; the depicted significance levels result from pairwise post-hoc tests versus the control. **(a,b)** Hippocampus. Anti-AT8: p = 0.163; anti-S404: p = 0.022; anti-S396: p = 0.574. **(c,d)** Cortex. Anti-AT8: p = 0.003; anti-S404: p = 0.016; anti-S396: p = 0.134. **(e)** Thalamus. Anti-AT8: p = 0.935; anti-S404: p = 0.353; anti-S396: p = 0.126.

APP expression in these mice (Fig. 4) was significantly induced in the hippocampus (p = 0.001) and modulated in the cortex (p = 0.007), whereas the thalamus showed a loss of expression (p = 0.041). Analysis of detergent-soluble Aβ levels yielded significant effects in terms of a temporally reversible induction of Aβ_40_in the hippocampus (p = 0.029) and the retrosplenial cortex (p = 0.016) (Fig. 4e). Aβ_42_ was reversibly induced only in the thalamus (p < 0.001), while the hippocampus and the cortex exhibited a late reduction of Aβ_42_ (p = 0.010 for hippocampus, p = 0.034 for cortex) (Fig. 4f). Total Aβ levels calculated per mouse and tissue as sum of Aβ_40_ and Aβ_42_ are shown in Fig. 4g and were significantly modulated in the hippocampus (p = 0.015), the thalamus (p = 0.014), the retrosplenial cortex (p = 0.019) and the caudate/putamen (p = 0.044). The general picture of these analyses is that Aβ was only induced in certain tissues and exclusively after 4 h, while MK-801 was actually present in the tissue, whereas elimination of the drug resulted in a return to baseline levels or even lower levels than before. The rapid elevation and subsequent drop of soluble Aβ is consistent with its short half-life of less than 2 h in vivo (Abramowski et al., 2008). Moreover, it concurs with seminal findings that synaptic activity at NMDA receptors regulated Aβ secretion into the interstitial fluid in APPswe/PS1 mice (Verges et al., 2011) and inhibited amyloidogenic APP processing in cortical cell culture (Hoey et al., 2009).

**Figure 4.**
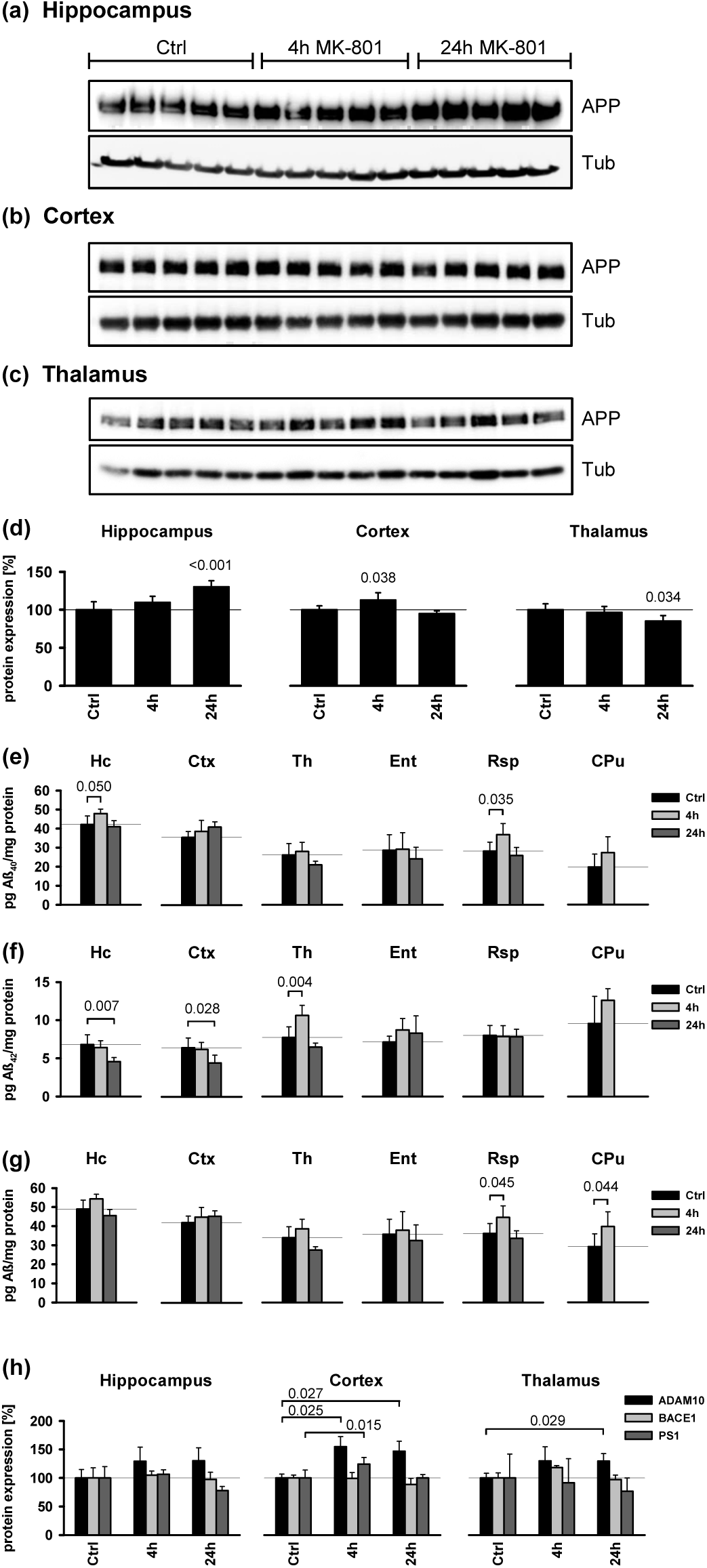
APP expression and processing in MK-801 treated mice. **(a-c)** APP immunoblots of three different brain regions from MK-801 treated mice. **(d)** Quantification of APP expression. Statistical analysis was done by One-way ANOVA as in Fig. 3. Hippocampus: p = 0.001, n = 5; cortex: p = 0.007, n = 5; thalamus: p = 0.041, n = 5. **(e)** Soluble SDS-tissue concentrations of Aβ_40_ (pg/mg protein) after 4 h and 24 h MK-801 treatment were determined by ELISA in the following brain regions:hippocampus (Hc): p = 0.029; cortex (Ctx): p = 0.136; thalamus (Th): p = 0.111; entorhinal cortex (Ent): p = 0.597; retrosplenial cortex (Rsp): p = 0.016; caudate/putamen (CPu): p = 0.149 by One-way ANOVA. In this assay, a control group of n = 7 animals was analyzed; treatment groups were n = 5 animals as in (a)-(d). The 24 h CPu tissue was not analyzed. **(f)** Concentrations of Aβ_42_ (pg/mg protein) in the same tissues as in (e). Hippocampus: p = 0.010; cortex: p = 0.034; thalamus: p < 0.001; entorhinal cortex: p = 0.266; retrosplenial cortex: p = 0.966; caudate/putamen: p = 0.135 by One-way ANOVA. **(g)** Total Aβ concentrations (pg/mg protein) calculated per mouse from the above Aβ_40_ and Aβ_42_ contents. Hippocampus: p = 0.015; cortex: p = 0.340; thalamus: p = 0.014; entorhinal cortex: p = 0.661; retrosplenial cortex: p = 0.019; caudate/putamen: p = 0.044. **(h)** Effect of MK-801 treatment on the expression of different secretases involved in APP processing as determined by Western blotting. ANOVA significance levels were: ADAM10 (hippocampus: p = 0.114; cortex: p = 0.026; thalamus: p = 0.043), BACE1 (hippocampus: p = 0.722; cortex: p = 0.171; thalamus: p = 0.218), PS1 (hippocampus: p = 0.023; cortex: p = 0.010; thalamus: p = 0.671). Statistical evaluations were done on n = 5 animals as in (d). The numbers in the graph represent pairwise significances from post-hoc tests. Actin was used as loading control on BACE1 blots.

Remarkably, the reversible excitatory blockade was apparently entrained in certain tissues in terms of a long-term induction of APP (especially in the hippocampus) and AT8 (especially in the cortex). A similar phenomenon of a rapid and persistent induction of cortical APP synthesis after loss of activating subcortical innervation has long been noted (Wallace et al., 1993). These observations are puzzling since APP does not tend to accumulate, but rather exhibits a short half-life (primary culture intracellular pool: ∼0.5 h; surface pool: ∼4 h) (Storey et al., 1999). What may be the functional role of the prolonged induction of APP?

We tested the hypothesis that proteolytic fragments other than Aβ may account for the long-term induction of APP after transient blockade of excitatory neurotransmission. Therefore, the expression of the main APP secretase proteins ADAM10, BACE1 and PS1 was analyzed by Western blotting (Fig. 4h). BACE1 was unregulated in the three investigated tissues. In contrast, the Aβ-producing γ-secretase subunit PS1 was reversibly induced in the cortex (p = 0.010) and belatedly decreased in the hippocampus (p = 0.023), which indicates that Aβ’s physiological role is to respond to excitatory insufficiency acutely, but not chronically. This interpretation is supported by the observation that ADAM10, an Aβ-preventing α-secretase, was persistently induced in the cortex (p = 0.026) and the thalamus (p = 0.043). The ensuing formation of the potent synaptotrophic effector sAPPα (Moechars et al., 1996; Bell et al., 2008; Nicolas and Hassan, 2014) or the known glutamate metabolism enhancer and synaptic regulator AICD (Schrenk-Siemens et al., 2008; Bukhari et al., 2017) may be the purpose of this induction, to permanently resolve the excitatory blockade.

The observed tissue-specific responses of tau and APP following NMDA receptor blockade resemble patterns seen in AD. The inertia of the thalamus to increase tau phosphorylation (Fig. 3e) and its selective induction of Aβ_42_ formation (Fig. 4f) correlate with the fact the thalamus in AD shows amyloid pathology very early, in fact as early as the hippocampus, but essentially lacks neurofibrillary tangles until very late in the disease (Aggleton et al., 2016). Moreover, the anterior thalamus is functionally interdependent with the retrosplenial cortex, one of the first areas affected in prodromal AD (Nestor et al., 2003). The impairment of both regions is thought to account for the early loss of episodic memory in the disease (Aggleton et al., 2016). The retrosplenial cortex also was the most responsive region in terms of Aβ_40_ induction in the current study (Fig. 4e).

In physiology and biochemistry, adaptive responses to pulsatile challenges often become weaker due to habituation. In the present paradigm, drug elimination may become accelerated, or the brain may reinforce its general capacity to endure NMDA receptor blockade, potentially by means unrelated to the AD-related molecules examined here. To investigate whether the AD-related molecules were indeed limiting and relevant after prolonged expose, the effect of repeated, three-times-weekly injections of MK-801 (1.0 mg/kg) over 6 weeks was studied in male and female wild-type mice. Analysis was done approximately 24 h after the last injection (∼10 half-lives), to exclusively detect changes persisting in the absence of the drug. In fact, we did not find significant differences between the treated group and the control in male mice treated for 6 weeks. Specifically, the 24 h elevations in hippocampal APP and S404 phosphorylation as well as cortical AT8 and S404 phosphorylation (Fig. 3b,d; Fig 4d) were absent (data not shown). Thus, ∼16-week-old male mice that had been treated with MK-801 for 6 weeks appeared to be capable of full habituation. With prolonged duration of treatment, however, habituation capacity started to decline, since mice treated with MK-801 for 12 weeks (and analyzed at an age of ∼22 weeks) displayed increased cortical AT8 (p = 0.039 by One-way ANOVA) and S404 (p = 0.011) phosphorylation again (Fig. 5c,d); APP expression in these mice was yet unchanged in the hippocampus and significantly lowered in the cortex (p = 0.039) (Fig. 5e).

**Figure 5.**
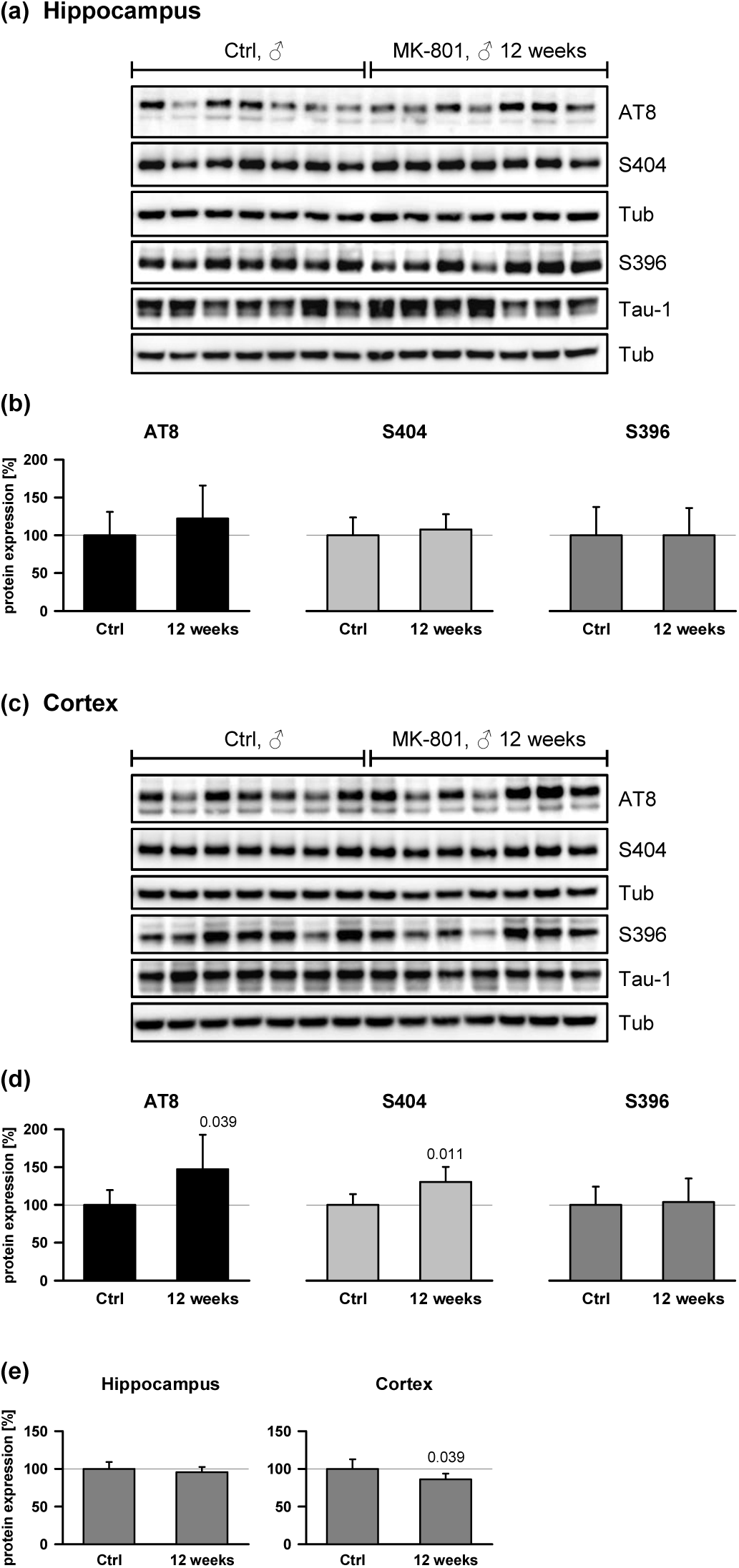
Effect of repeated, 12-week MK-801 treatment in male mice. Male mice were treated with MK-801 (1.0 mg/kg ip) three times per week (Mon, Wed, Fri) for 12 weeks (36 injections altogether). Tissue sampling was performed 24 h after the last treatment, analyzing the hippocampus and the total cortex (compare the Methods) for tau phosphorylation and APP expression. Statistical evaluations were done by One-way ANOVA. **(a)** Hippocampus immunoblots analyzing saline-treated control animals (n = 7) and the treatment group (n = 7) for specific tau phosphorylation. **(b)** Quantified hippocampal expression levels detected with antibodies anti-AT8: p = 0.335, anti-S404: p = 0.554, anti-S396: p = 0.996. Signals were Tub-normalized and related to total tau expression. **(c)** The same analysis as in (a) done for the cortex. **(d)** Quantified expression levels in the cortex. Anti-AT8: p = 0.039; anti-S404: p = 0.011; anti-S396: p = 0.819. **(e)** Quantification of APP expression after repeated MK-801 treatment (n = 7 animals per group). Hippocampus: p = 0.362; cortex: p = 0.039.

In contrast to male mice, female mice treated under the 6-week-paradigm did demonstrate significantly induced AT8 phosphorylation in the hippocampus (p = 0.004) (Fig. 6a,b) as well as S404 phosphorylation in the hippocampus (p = 0.004) and the cortex (p = 0.002) (Fig. 6c,d). APP expression levels were unchanged in both regions (Fig. 6e), whereas a significant loss of PS1 expression was observed (Fig. 6f), concurring with the results from single-treatment male mice (Fig. 4h). To explore the robustness of these effects, a further group of female mice treated with a lower dose of MK-801 (0.2 mg/kg) for 6 weeks was studied. Analysis of tau phosphorylation in the cerebral cortex (Fig. 7) indicated significantly increased levels of all three investigated epitopes, AT8 (p = 0.002), S404 (p = 0.004) and S396 (p = 0.009).

**Figure 6.**
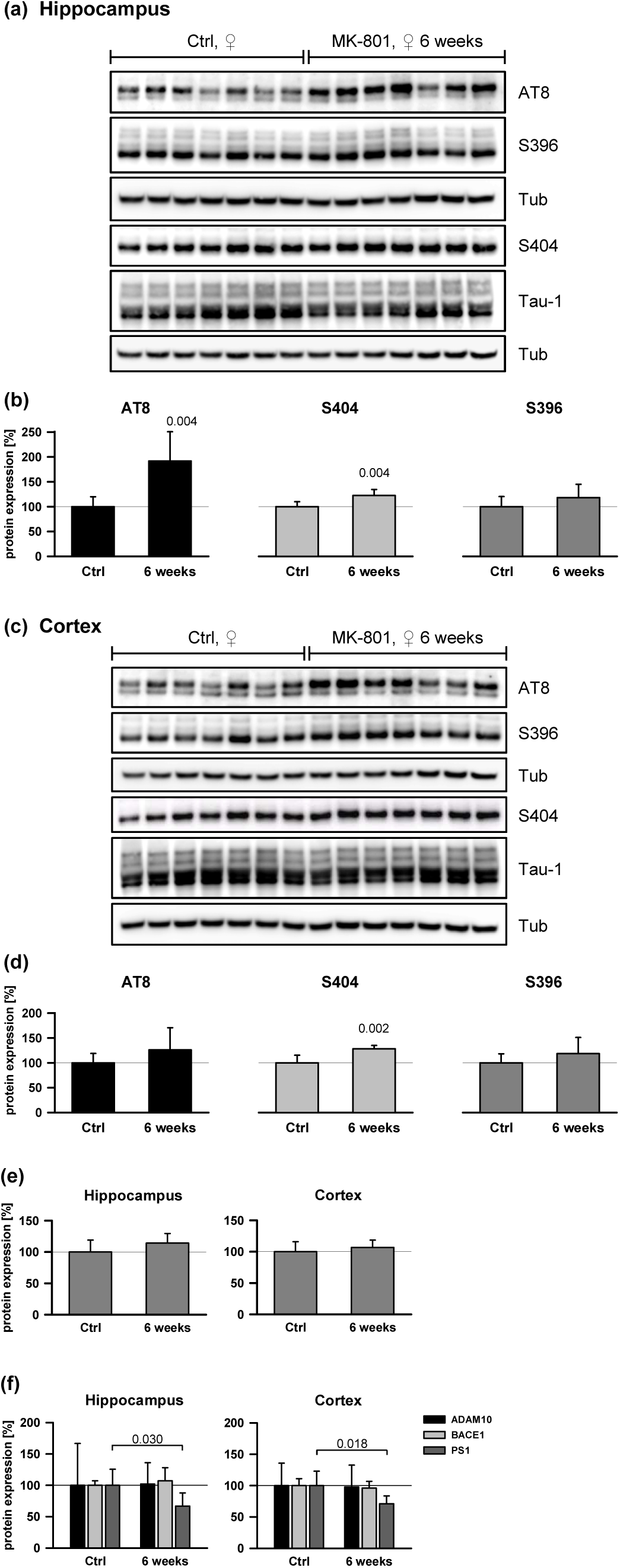
Effect of repeated MK-801 treatment in female mice. Female mice were treated with MK-801 (1.0 mg/kg ip) three times per week for 6 weeks (18 injections). Tissue sampling was performed 24 h after the last treatment. **(a)** Hippocampus immunoblots analyzing tau phosphorylation. **(b)** Quantified hippocampal phospho-tau expression detected with antibodies anti-AT8: p = 0.004, anti-S404: p = 0.004, anti-S396: p = 0.201. **(c)** The same analysis as in (a) done for the cortex. **(d).**Quantified expression levels in the cortex. Anti-AT8: p = 0.216; anti-S404: p = 0.002; anti-S396: p = 0.252. **(e)** Quantification of APP expression (n = 7 animals per group). Hippocampus: p = 0.176; cortex: p = 0.418. **(f)** Expression of APP-processing secretases. ANOVA significance levels were: ADAM10 (hippocampus: p = 0.949; cortex: p = 0.196), BACE1 (hippocampus: p = 0.434; cortex: p = 0.552), PS1 (hippocampus: p = 0.030; cortex: p = 0.018). Statistical evaluations were done on n = 7 animals as in (e). Actin was used as loading control for BACE1.

**Figure 7.**
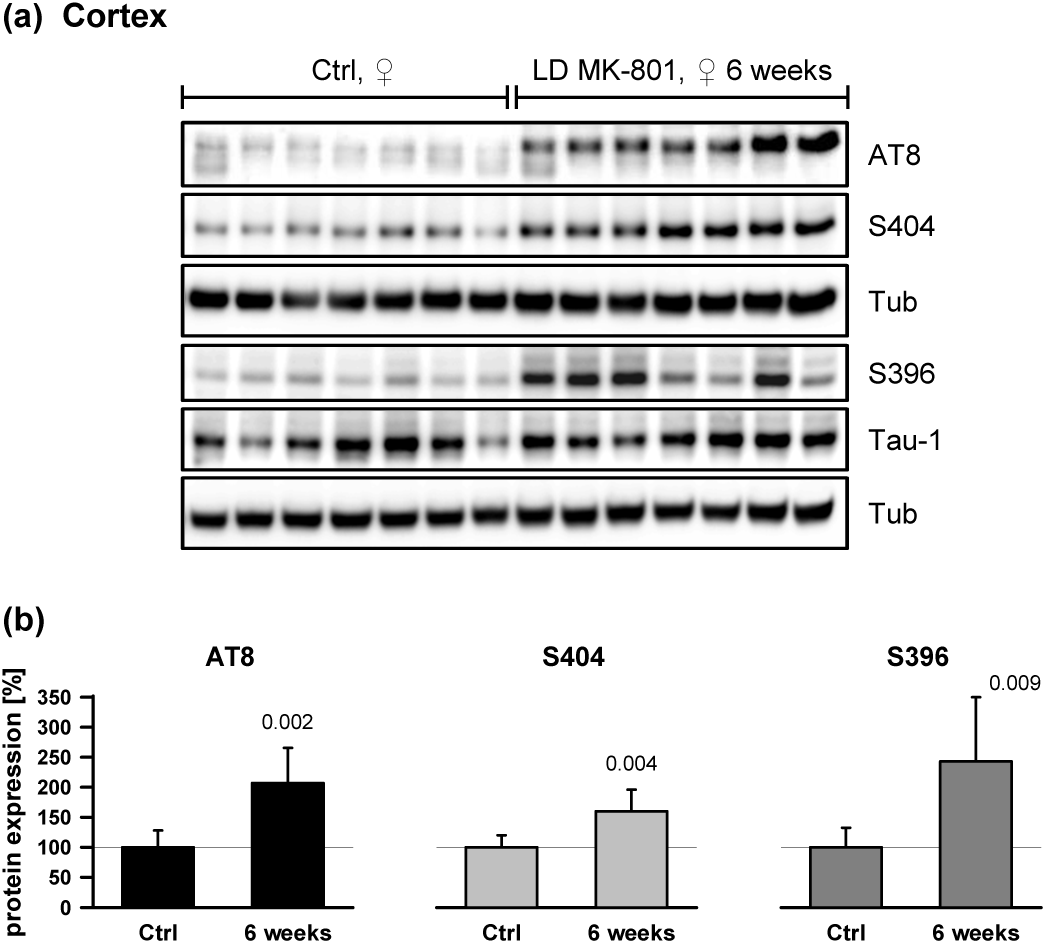
Effect of repeated low-dose MK-801 treatment in female mice. Female mice were treated with low-dose (LD) MK-801 (0.2 mg/kg ip) three times per week for 6 weeks as in the normal-dose experiments in Fig. 6. Tissue sampling was performed 24 h after the last treatment. **(a)** Cortical samples from control animals (n = 7) and the treatment group (n = 7) were assayed for specific tau phosphorylation. **(b)** Quantification of tau phosphorylation as detected with antibodies anti-AT8: p = 0.002, anti-S404: p = 0.004, anti-S396: p = 0.009. Signals were Tub-normalized and afterwards related to total tau expression.

The notable difference of male and female mice in this study parallels AD epidemiology (Snyder et al., 2016) and may be related to the established, higher sensitivity of females to toxic NMDA receptor inhibition, which has been shown to be critically influenced by gonadal hormone levels (De Olmos et al., 2008). More broadly, these hormones are known to have a strong impact on hippocampal functioning (Spencer et al., 2008). Alternatively, but not mutually exclusive, the weaker habituation in females might be related to the relatively higher brain levels of MK-801 after ip injection (Andine et al., 1999). In summary, the data in Fig. 5-7 indicate that repeated NMDA receptor blockade bears the potential to induce in vivo, in both sexes and beyond the acute stage, relevant markers of AD pathology such as cortical AT8 and S404 immunoreactivity (Mondragon-Rodriguez et al., 2014).

## Discussion

### A chronic NMDA receptor insufficiency model for Alzheimer’s disease

AD displays a distinctive histopathology foremost characterized by neurofibrillary tangles and amyloid plaques (Braak et al., 1994; Roher et al., 2014). Both these features have been viewed to be causal for the disease. We propose that both alterations reflect the culmination of a derailed long-lasting pursuit of postsynaptic neurons to receive sufficient excitatory input as calibrated during early development (Fig. 8). According to this model, excitatory insufficiency is caused by functional inhibition of excitatory neurotransmission in sporadic AD (e.g. due to the presence of ApoE4 protein or anti-NMDA receptor antibodies), and by genetically determined hypogenesis of excitatory structures like fiber tracts in familial AD (see below for references). Both variants would result in the same compensatory, but unfortunately perpetuated response of induced Aβ formation and tau phosphorylation that antecedes plaque and tangle formation.

**Figure 8.**
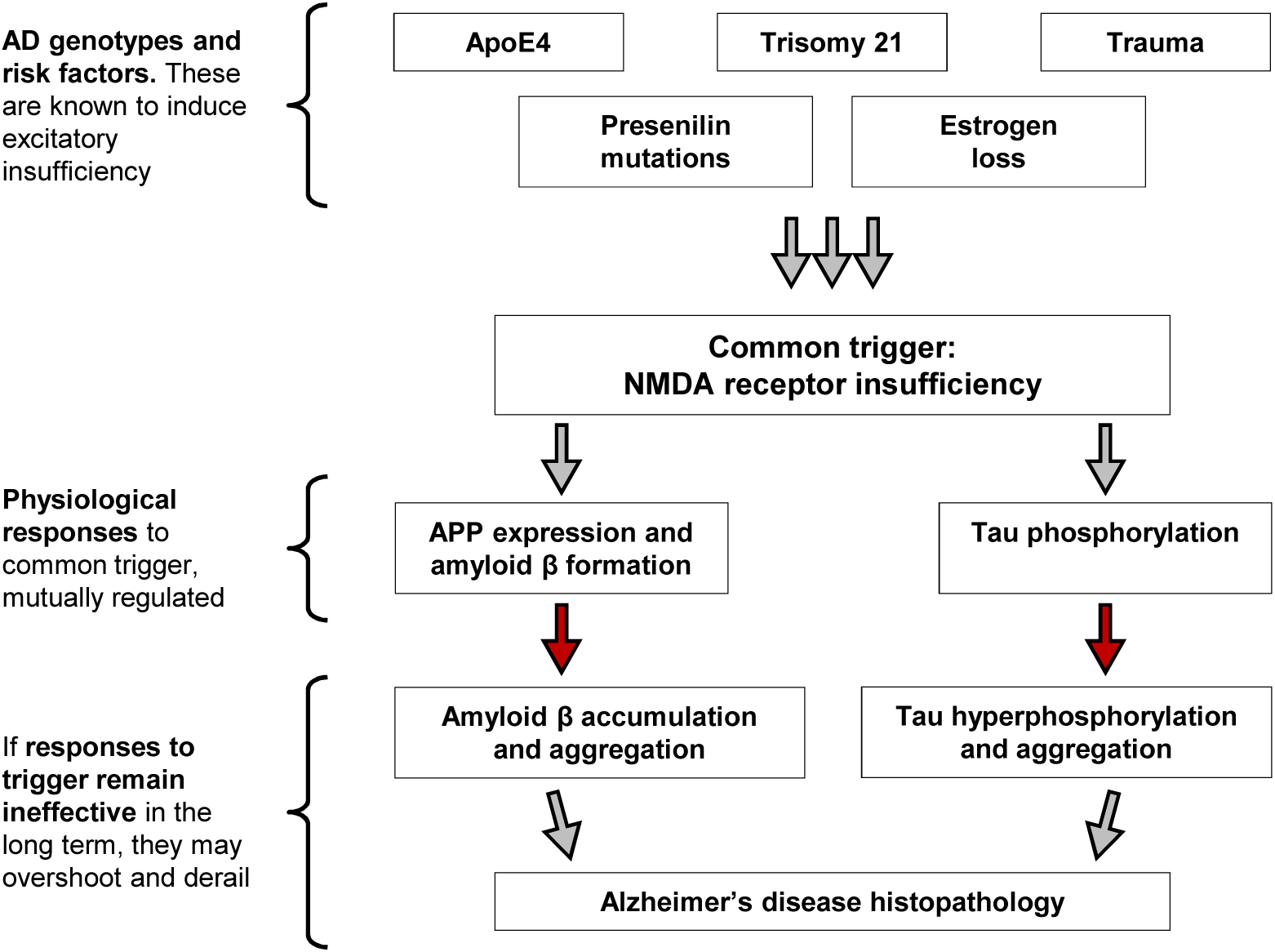
Proposed model for the etiology of AD. According to this model, the common pathogenetic mechanism of AD is insufficient excitatory neurotransmission via NMDA receptors. The insufficiency is caused by genetically determined, anatomically manifest hypogenesis of excitatory structures in familial AD and trisomy 21-related AD, by intracellular trapping of synaptic NMDA receptors in ApoE4-related AD, and by NMDA antagonistic effectors (traumata, autoantibodies) in sporadic AD. In all cases, a regulatory response of initially elevated APP expression, Aβ formation and tau phosphorylation is induced by the insufficiency, as shown in this study using pharmacological NMDA antagonists. The evoked effects are directed at the interim stabilization and permanent resurrection of excitatory neurotransmission after temporary lesions, as referenced in the Discussion. Since none of the induced response molecules are capable of annihilating sources of chronic NMDA receptor insufficiency (e.g. none can remove autoantibodies), the triggered regulatory responses become chronic and dysregulated themselves, resulting in tau hyperphosphorylation and aberrantly elevated Aβ levels. In the end, both molecules aggregate and form visible lesions.

Functionally, it is inferred that Aβ serves as acute glutamatergic sensitizer (Mattson et al., 1992; Harkany et al., 2000; Puzzo et al., 2008; Garcia-Osta and Alberini, 2009; Morley et al., 2010; Danysz and Parsons, 2012) intended only for short-term use, which is rapidly replaced by other APP cleavage products such as sAPPα to serve as synaptotrophic factors (Moechars et al., 1996; Bell et al., 2008; Nicolas and Hassan, 2014). Tau phosphorylation serves a similar purpose by either facilitating synaptic protein transport (Dixit et al., 2008), sensitizing NMDA receptors (Ittner et al., 2010), or both. Synapse fortification or synaptogenesis (as per sAPPα) are superior to synaptic sensitization (as per Aβ) since simple sensitization bears the risk of self-excitation (seizures) and does not increase the signal-to-noise ratio of synaptic transmission, whereas synapse formation genuinely does. Notably, diminished excitatory signal-to-noise ratios appear to be common in AD (Danysz and Parsons, 2012) and may underlie the considerably increased risk of seizures especially in familial forms of the disease (Pandis and Scarmeas, 2012). However, synapse formation requires time and coordination, why the interim time is bridged by default with a glutamatergic sensitizer to maintain functional integrity in the short-term. This idea is supported by our findings that Aβ was only transiently elevated after NMDA antagonist treatment, without any induction of BACE1, with only temporary induction of PS1 in the cortex and a late reduction of PS1 expression in the hippocampus (Fig. 4h). A persistent reduction of PS1 was also seen in female mice after subchronic NMDA antagonist treatment (Fig. 6f), which might functionally recapitulate the inhibitory effects of AD-related PS1 mutations (Shen and Kelleher, 2007; Xia et al., 2015; Sun et al., 2017). In contrast, ADAM10 was robustly upregulated in the cortex 4 h and 24 h after single antagonist injection (Fig. 4h). Similar observations of increased ADAM10, but unchanged BACE and reduced PS1 expression have been made in sporadic AD tissues (Takami et al., 1997; Davidsson et al., 2001; Gatta et al., 2002). Thus, APP processing yields at least two products, Aβ and sAPPα, which shall remedy adverse consequences of neuronal deprivation from excitation. In endocrinology, such complementary regulatory functions of proteolytic fragments coming from the same precursor protein are quite common, as in angiotensinogen (Santos et al., 2013) or proopiomelanocortin (O’Donohue and Dorsa, 1982).

The “compensatory response” model of AD provides an explanation why many genetic and sporadic triggers of AD either disrupt the development of glutamatergic connections during infancy (e.g. PS1 and APP mutations) or adult glutamatergic signaling (e.g. ApoE4 alleles, repeated mechanical injuries), as discussed in the following.

### Explanatory scope of the proposed model

#### I. AD occurs in hereditary and sporadic forms

A small fraction of AD cases are caused by hereditary mutations in APP, PS1 and PS2, or by gene duplications in the region of the APP gene. Traditionally, these genetic alterations have been viewed to be associated with persistently elevated levels of Aβ or particularly toxic subspecies thereof (Hardy and Selkoe, 2002). Such cases may be easily accommodated into the excitatory insufficiency model by considering brain development as follows. Glutamatergic tone is very strictly regulated during neuronal development (Spitzer, 2006) since hypoactivity of the glutamatergic system leads to immediate neuronal death (Ikonomidou et al., 1999), whereas hyperactivity might lead to convulsions, epilepsy and other life-threatening sequelae (Chapman, 1998; Coulter and Eid, 2013). Thus, elevated levels of the glutamatergic sensitizer Aβ during infancy will induce a compensatory, irreversible decrease in the baseline structural components of excitatory neurotransmission, such as the number of axons in excitatory fiber tracts or the number of neurons. These reduced structures will be affected much earlier by any form of subtle, adult or age-related excitatory insufficiency as they are (i) fewer and (ii) already run at a high rate of Aβ production to maintain baseline neurotransmission. Hence, the early and regular pathology in the hereditary forms of AD could in fact be attributable to developmental excitatory insufficiency on a structural level.

This notion is supported by an extensive body of evidence. For instance, temporally limited induction of Aβ production during early postnatal development (in on/off APPswe/ind mice) has been shown to result in permanent locomotor hyperactivity in adult animals (Rodgers et al., 2012), which is a typical phenotype of NMDA antagonist exposure (Andine et al., 1999). Significantly reduced numbers of presynaptic terminals and neurons have been counted in the hippocampus of young, 2-3 month-old APPind mice (Hsia et al., 1999), together with a shortened corpus callosum and a smaller dentate gyrus by 100 days of age (Redwine et al., 2003). Reduced spine density in this region high in NMDA receptors has also been observed in young adult, 4 month-old APPswe mice (Jacobsen et al., 2006). In humans, heterozygote ApoE4 carriers have been described to exhibit entorhinal cortex thinning (Shaw et al., 2007) and grid cell dysfunction (Kunz et al., 2015) as soon as in their early 20s, long before their increased risk of entorhinal pathology and AD precipitates. Grid cells are selectively dependent on NMDA receptors for firing (Gil et al., 2018). Brain structural and functional abnormalities have also been reported for young adult carriers of autosomal-dominant PS1 mutations (Reiman et al., 2012), including the need for hippocampal metabolic hyperactivation to achieve the same results as non-carriers in a learning task (Quiroz et al., 2010). Furthermore, unbalanced inhibitory activity resulting from neurodevelopmental defects appears to be a central component of Down syndrome (Ferrer and Gullotta, 1990; Belichenko et al., 2004; Fernandez et al., 2007), which later in life is characterized by a greatly increased risk of AD (Wiseman et al., 2015). Finally, several other proteins involved in neuronal development have emerged from sporadic AD GWAS studies, such as MEF2C, EPHA1, and SLC24A4 (Lambert et al., 2013; Rosenthal et al., 2014). Thus, carriers of AD mutations or risk alleles appear to be anatomically different and hypo-excitatory (with metabolic and Aβ compensation) already during brain development.

Less traditionally, the idea has emerged that mutations in PS1, underlying the vast majority of familial AD cases, may not result in any systematically elevated or uniformly shifted production of Aβ species (Sun et al., 2017), but may rather evoke immediate neurodevelopmental defects and excitatory network failure via a PS1 loss of function (Shen and Kelleher, 2007; Xia et al., 2015). Notably, these developmental defects compromise functions that critically depend on glutamatergic integrity such as LTP (Xia et al., 2015). Comparable observations of early impaired LTP have been made in mutant APP transgenic mice (Hsia et al., 1999; Moechars et al., 1999; Jacobsen et al., 2006). Moreover, an explicit NMDA receptor deficit has been described for PS1/2 knockout mice (Saura et al., 2004). Hence, under the premises of the “less traditional” interpretation, familial AD would immediately fit into an NMDA receptor insufficiency model of AD causation.

#### II. Amyloid and tau pathology are common in many CNS disorders beyond AD

Amyloid plaques as well as neurofibrillary tangles are widespread and occur isolated or in combination in many different nosological entities, such as frontotemporal dementia (Gordon et al., 2016), cerebral amyloid angiopathy (Revesz et al., 2002) or Down syndrome (Wiseman et al., 2015). In addition, clinically diagnosed AD patients are not infrequently “amyloid negative” (Monsell et al., 2015; Landau et al., 2016), while cognitively normal patients can harbor extensive amyloid pathology (Elobeid et al., 2014). Consequently, it appears that both lesions are rather general phenomena in the brain that occur in association with rather diverse problems of the brain. General occurrence of a specific molecular phenotype is yet much better reconcilable with a “compensatory response” model than a “disease cause” model, as outlined on the following two examples.

a. Acute CNS lesions. Induction of APP has long been known to constitute a rapid and persistent response of the cortex to deficient subcortical innervation (Wallace et al., 1993). Notably, this effect was seen after inhibition of cholinergic as well as adrenergic and serotonergic pathways in rats (through chemical lesions in the nucleus basalis of Meynert, the ascending noradrenergic bundle and the dorsal raphe nucleus, respectively). Formation of Aβ as one consequence of such APP induction has been widely confirmed in experimental traumatic brain injury (Bird et al., 2016). Related findings have been made in humans, where APP has been used as early marker of axonal damage after mechanical trauma (Sherriff et al., 1994). Severe neurotrauma patients within hours can develop diffuse amyloid plaques (Ikonomovic et al., 2004) whose long-term persistence is likely (Scott et al., 2016). By correlation, higher early Aβ levels have been found to predict better clinical outcome after severe neurotrauma (Magnoni et al., 2012). More subtle and specific inhibition of axonal transport by kinesin reduction has also been shown to enhance Aβ production in APPswe mice (Stokin et al., 2005), whereas the inhibition of the Aβ response was associated with poorer recovery after spinal cord injury in wild-type mice (Pajoohesh-Ganji et al., 2014). Readily applicable to all these cases is the idea that APP expression and Aβ formation occurred as compensatory response to the lesional excitatory interruption and were directed at a provisional functional maintenance of the lesioned site, leading the way for its later repair.
b. Down syndrome (DS). DS (or trisomy 21) is strongly associated with dementia and AD-like pathology at a relatively early age (Wiseman et al., 2015). It has been proposed that the increased gene dose of APP (which is encoded on chromosome 21) might be causative to this observation, by directly leading to an increased production of APP and Aβ (Rumble et al., 1989). However, a consistent induction of APP over ages and cell types does not occur in DS, calling for specific regulatory mechanisms to induce pathological amyloid formation (Antonarakis, 2017). In addition, overexpression of wild-type APP alone is not related to neurodegeneration or plaque pathology in mice (Elder et al., 2010). Hence, it is more plausible to conclude that excitatory network failure of neurodevelopmental origin in DS causes a persistent compensatory elevation of the glutamatergic sensitizer Aβ, resulting in untimely amyloid pathology. In fact, DS has been associated with selectively suppressed excitatory (Belichenko et al., 2004; Kurt et al., 2000) and excessive inhibitory activity (Fernandez et al., 2007) as well as an increased risk of seizures due to an exaggerated amplification of small excitatory signals in an overly inhibitory background (Araujo et al., 2015).

#### III. ApoE polymorphisms are predominant risk factors for sporadic AD

The strong association of ApoE4 with sporadic AD (Rosenthal and Kamboh, 2014; Deming et al., 2017) first reported in 1993 (Strittmatter et al., 1993) has entailed numerous, often contradictory hypotheses on the biochemical origin of this association, ranging from cholesterol transport to Aβ clearance (Liu et al., 2013). Still, there is one mechanism that may be viewed to stand out simply for its effect size and its localization to the synapse, the prime site of AD (Terry et al., 1991). After binding to its postsynaptic receptor ApoER2, ApoE is internalized, followed by eventual dissociation and recycling of ApoER2 to the surface. Notably, ApoE4 remains internalized much longer than other isoforms (Heeren et al., 2004), leading to impaired surface recycling of ApoER2 and co-endocytosed NMDA receptors (Chen et al., 2010). In practice, ApoE4 has been shown to reduce neuronal NMDA receptor surface density by a factor of 3-5 compared to ApoE3 and ApoE2 under relevant in vitro conditions (Chen et al., 2010). Furthermore, ApoER2 is a canonical receptor for reelin, a pivotal synaptotrophic effector (Tissir and Goffinet, 2003) and activator of NMDA receptors via Src kinase-mediated phosphorylation (Beffert et al., 2005). After competitive ligation of ApoER2 with ApoE4, but not with ApoE3 or ApoE2, reelin’s capacity to enhance long-term potentiation was virtually abolished (Chen et al., 2010). Therefore, ApoE4 is a specific and severe NMDA receptor functional suppressor at the postsynaptic membrane. Accordingly, undue amplification of weak excitatory signals has been reported for ApoE4 knock-in mice (Nuriel et al., 2017). This compensatory phenotype is likely related to the substantially elevated Aβ levels shared by ApoE4 knock-in mice (Liraz et al., 2013), ApoE4/APPind mice (Bales et al., 2009) and human ApoE4 carriers (Monsell et al., 2015).

#### IV. Association of AD with sex, sleep, and auditory performance

Epidemiology and clinic have yielded a variety of apparently unrelated associations of AD with specific predictive factors. We find that many of these become plausible after introducing NMDA receptor insufficiency as disease trigger. We briefly discuss three of them below.

i. Sex. Amyloid pathology in humans begins at about 50 years of age for both sexes, but progresses significantly faster in females (Braak et al., 2011). This meshes with the fact that NMDA receptor density and function in the hippocampus are strongly augmented by estrogen (Wooley et al., 1997; Spencer et al., 2008), which drops in females after the age of 50 years. Aggravated NMDA receptor insufficiency due to estrogen loss would thus require a more pronounced compensatory amyloid response in females. In fact, an inverse correlation of amyloid formation and estrogen levels in AD animal models and human patients has been noted (Li et al., 2014).
ii. Sleep. Recent evidence from a series of knock-out mice indicates that Ca^2+^-dependent hyperpolarization, which is strongly dependent on NMDA receptors, critically regulates sleep duration (Tatsuki et al., 2016). Correspondingly, sleep duration in animals was impaired by pharmacological NMDA receptor blockade (Tatsuki et al., 2016). Sleep disturbances, especially fragmented sleep, are recurrently seen in early AD (Bubu et al., 2017), which might thus reflect emerging NMDA receptor insufficiency.
iii. Auditory performance. Impaired hearing is one of the most significant correlates of dementia (Livingston et al., 2017) and very frequent in DS (Kreicher et al., 2018). Even if auditory functioning depends on various systems, it is noteworthy that subtle calcium dysregulation as per NMDA receptor inhibition seems to be predestined to cause auditory malfunction. For instance, the cochlea, the colliculus and the auditory cortex all express functional NMDA receptors that have been shown to be indispensable for auditory space map formation (Schnupp et al., 1995) and auditosensory processing (Bickel et al., 2008), but also for the development of noise-induced hearing loss (Sanchez et al., 2015). Moreover, hearing loss is the prime phenotype of mice with mutant PMCAs (Bortolozzi and Mammano, 2018), which are calcium pumps that in close cooperation with NMDA receptors determine the shape of neuronal calcium transients (Scheuss et al., 2006). In fact, deficient auditory plasticity has been shown to be amenable to treatment with NMDA agonists in the clinic (Kantrowitz et al., 2016).

### What might be the source of NMDA receptor insufficiency in sporadic Alzheimer’s disease?

i. Autoimmune antibodies. Anti-NMDA receptor autoantibodies are frequent (seroprevalence 10-20%, rising with age) (Zerche et al., 2015), act anti-excitatory in vitro (Pruss et al., 2012; Castillo-Gomez et al., 2016) and appear to be broadly associated with cognitive deficits and dementia (Pruss et al., 2012; Doss et al., 2014; Finke et al., 2017). Notably, their association with outcome in clinical stroke depends on the ApoE genotype (Zerche et al., 2015). Other autoantibodies against excitatory neurotransmitter receptors exist (Borroni et al., 2017). A role of autoantibodies in AD could be reconcilable with the recurrent appearance of microglial immune-response genes in AD GWAS studies (Sims et al., 2017), as microglia are the predominant antigen-presenting cells in the CNS (Aloisi et al., 2000).
ii. Excitatory network failure. Beyond genetic reasons as in DS or in ApoE4 carriers, acquired and persistent excitatory insufficiency may result from many presumed AD risk factors such as mechanical lesions (Li et al., 2017), periodic hemorrhages (Revesz et al., 2002; Cordonnier and van der Flier, 2011), or neurotropic viral infections (Itzhaki et al., 2016). Such events could also induce autoantibody formation as a secondary effect.

In summary, we have shown here that pharmacological blockade of NMDA receptors under physiological and non-transgenic conditions elicits a rapid regulatory response in terms of a tissue-specific induction of APP expression, Aβ formation and APP secretase modulation (Fig. 4) as well as tau phosphorylation (Fig. 3) in vivo. Very similar effects were seen under fully defined conditions of cortical cell culture in vitro (Fig. 1 and 2). Especially female mice were unable to biochemically habituate to repeated injections of the employed NMDA receptor blocker, indicating the pathogenic potential of long-term NMDA receptor insufficiency (Fig. 5-7). The derived hypothesis that persistent NMDA receptor insufficiency was the mechanistic cause of AD might open viable new means of prevention, i.e. the removal of circulating autoantibodies or the positive modulation of glutamatergic neurotransmission. Approved or repurposable drugs for both indications exist. These new approaches might complement more conventional measures such as the avoidance of brain injury or the continued stimulation of the brain through sensory, cognitive and motor challenges (Barnes and Yaffe, 2011), whose probable efficacy might also rely on ameliorated connectivity via NMDA receptors.

## Acknowledgements

We thank Roland Rabl, Annamaria Molnar, Martina Farcher, Manuela Prokesch, Petra Baumgartner, Stefanie Flunkert, Klaus Lorenzoni, and Birgit Hutter-Paier, all affiliated with QPS Austria, for their excellent management and professional realization of the animal studies. The funding of this work by the Volkswagen-Stiftung (Grants #87736, #89854 and #93286) is gratefully acknowledged.

## Author contributions

Both authors designed the experiments, the first author performed the experiments, both authors analyzed the data and wrote the paper.

## Methods

### Primary cortical cell culture

Primary cortical neurons were prepared from E17 Spraque-Dawley rats (Janvier Labs, Le Genest-Saint-Isle, France). The dams were anesthetized with isofluran and killed by guillotine before the progeny was removed. Embryos were decapitated, and cortices were prepared under a binocular microscope. Loosely chopped tissue pieces were then incubated in PBS containing 0.1% trypsin and 0.2% EDTA for 20 min at 37°C. To stop the trypsination, the tissue was transferred to Ca^2+^- and Mg^2+^-free HBSS supplemented with 10% FCS and carefully resuspended. After filtration through a sterilized 50 µm nylon gauze to remove undissociated tissue pieces, the cells were centrifuged at 1,200 g for 4 min at 22°C. The obtained pellet was resuspended in serum-free Neurobasal Medium supplemented with 1x B-27 supplement, 2 mM GlutaMAX and 25 µg/ml gentamicin. All buffers, media and supplements were from Thermo Fisher Scientific, Rockford, IL, USA. An aliquot of the cell suspension was mixed 1:1 with trypan blue solution (Sigma-Aldrich, St. Louis, MO, USA) before visually counting the intact cells in a hemocytometer. The cells were then seeded at a density of 80,000 cells/cm^2^ in 6-well, 24-well or 96-well plates (TPP, Trasadingen, Switzerland) that had been pre-coated for 30 min at RT with 0.1 mg/ml poly-L-ornithine (MW range 30 - 70 kDa, from Sigma-Aldrich). Cells were cultivated in the original, supplemented Neurobasal Medium in a humidified cell culture incubator under 5% CO_2_ at 37°C for 14 to 17 days.

For the experiments, the cultivation medium was replaced by fully defined physiological Ringer solution (125 mM NaCl, 2.5 mM KCl, 1.25 mM NaH_2_PO_4,_ 25 mM D-Glucose, 25 mM NaHCO_3,_ ultra-pure water) that had been pre-incubated in a cell culture incubator for 24 h to attain a pH of 7.4. Immediately afterwards, 2 mM CaCl_2_ was added manually to each well. Half of the subsequent experiments were done at baseline KCl concentrations of 2.5 mM (denoted “low” K+), the other half was done at higher and depolarizing KCl concentrations of 27.5 mM (denoted “high” K+) attained by adding 25 mM KCl to the respective cultures. Chemicals for the physiological Ringer solution were obtained at the highest available purity from Carl Roth, Karlsruhe, Germany. The cells were then treated with either of the following agents: 100 µM D-AP5, a highly specific NMDA receptor antagonist (Biotrend, Köln, Germany, or Cayman Chemical, Ann Arbor, MI, USA), 5 µM N-methyl-D-aspartate (NMDA) (Biotrend or Sigma-Aldrich), 5 µM L-glutamate (Sigma-Aldrich), or combinations thereof. All compounds had been dissolved as 100x stocks in ultra-pure water followed by sterile filtration. After addition of the compounds, the plates were incubated for 4 h before the incubation Ringer solution was removed and the cells were collected in lysis buffer (100 mM Tris, 20% sucrose, 0.5% SDS, pH 7.4 with HCl) supplemented with 1x protease inhibitor cocktail (P8340 from Sigma-Aldrich) and 1x phosphatase inhibitor cocktail (PhosSTOP from Roche, Mannheim, Germany). After brief sonication, the protein concentration of the samples was determined colorimetrically using bicinchoninic acid-copper (BCA Protein Assay Kit from Thermo Fisher Scientific). Samples were stored at -25°C.

### Cell survival and metabolic activity assays

Neuronal survival and cellular integrity in cell culture were monitored qualitatively at regular intervals using phase-contrast microscopy. Any cultures showing signs of beginning cellular disintegration, glial proliferation or floating debris were discarded. Quantitative assessment of neuronal survival was achieved by immunostaining of MAP-2-positive cells followed by cell counting (Hajieva et al., 2015). To this end, cells were cultivated on coated glass cover slips in 24-well plates and treated with glutamatergic agonists and antagonists as described before. Thereupon, the Ringer solution was replaced by -80°C methanol, and the cells were fixed for at least 1 h at -80°C. The methanol was removed, and the cells were washed with PBS and blocked for 1 h with 3% FCS in PBS at room temperature. Immediately after blocking, the cells were supplied with the primary antibody (anti-MAP-2, M4403, lot #035M4780V from Sigma-Aldrich, dilution 1:500 in blocking solution) and incubated overnight at 4°C. After three washes with PBS, the secondary antibody (Cy3-conjugated donkey anti-mouse IgG, 715-165-151, lot #123602 from Jackson Immunoresearch, West Grove, USA, diluted 1:200 in PBS) was applied. After 1 h incubation at room temperature in the dark, the secondary antibody was removed, and the cells were washed again. The nucleic acid dye 4′,6-diamidino-2-phenylindole (DAPI, from Sigma-Aldrich) was used as nuclear counterstain to correct for unequal plating and was applied at this stage as 1 µg/ml solution in PBS for 15 min. After three further washes with PBS, fluorescence images of the stained coverslips were taken. Three different pictures per coverslip from two coverslips per cell culture plate were counted to generate the data for one biological replicate. The added numbers of MAP-2-positive cells were then related to the added numbers of DAPI-positive nuclei, with untreated control cultures serving as 100% reference.

To quantify overall cell viability in terms of metabolic activity, MTT assays were conducted as described (Hajieva et al., 2015). Cells were cultivated in 96-well plates and treated as described before. After 3 h incubation with the agents of interest, the cells were supplied with 0.5 mg/ml MTT (Sigma-Aldrich) under unaltered experimental conditions. After 1 h, the reduction of MTT was stopped by adding 1 volume solubilization solution (40% dimethylformamide, 10% SDS, pH 4.0 with acetic acid). The plates were then incubated for 16-24 h in the dark to lyse the cells and the precipitated formazan crystals, before reading the plate at 560 nm. Results from 5 wells on one plate were averaged and related to the control to generate the data for one biological replicate.

### Animal studies

#### Single treatment study

Male C57BL/6JRj mice (Janvier Labs) at an age of 10 weeks were intraperitoneally (ip) injected with either saline (n = 7 animals) or 1.0 mg/kg (+)-MK-801 hydrogen maleate (M107, lot #080M4610V from Sigma-Aldrich) dissolved in saline (n = 10 animals). Four hours after the injection, the control group and n = 5 animals of the MK-801 treated animals were killed by cervical dislocation. The other n = 5 MK-801 treated animals were killed 24 h after compound injection by the same method. Whole brains were prepared and rapidly frozen in native state on dry ice before storage at -80°C.

#### Repeated treatment study

Male and female C57BL/6JRj mice (Janvier Labs) aged 10 to 12 weeks were repeatedly injected ip with saline or 1.0 mg/kg (+)-MK-801 hydrogen maleate (M107, lot #105M4606V from Sigma-Aldrich) dissolved in saline. An additional low-dose (LD) group of female mice was repeatedly injected ip with 0.2 mg/kg MK-801 in the same manner. The treatments occurred three times per week (Mon, Wed, Fri) at about the same time of day for a period of 6 or 12 weeks as indicated, totaling to a number of r = 18 or r = 36 injections, respectively. Groups sizes were n = 8 saline controls and n = 8 treated animals for 6-week males, n = 8 saline controls and n = 8 treated animals for 12-week males, and n = 8 saline controls, n = 8 treated animals and n = 8 low-dose treated animals for females (all treated for 6 weeks). Animals were sacrificed 24 h after the last injection by cervical dislocation as in the single treatment study. Mouse brains were prepared and snap frozen on dry ice before storage at -80°C.

#### Workflow and sample shipment

All animal experiments in mice and the preparation of whole brains were done by QPS Austria, Neuropharmacology group (Grambach, Austria). QPS Austria (part of QPS Holdings) is a full-service contract research organization (CRO) providing preclinical and clinical research services. Whole brains were shipped at -80°C from Grambach to Mainz by courier. Further tissue dissection and all analytical procedures were performed at the Institute for Pathobiochemistry of the University of Mainz, Germany.

#### Animal ethics

The QPS Austria animal facility is fully accredited by the Association for Assessment and Accreditation of Laboratory Animal Care (AAALAC). All procedures in the single treatment study and the repeated treatment study complied with the Animal Care and Welfare Committee. Animals were maintained according to the animal welfare regulations of the Ministry of Science of the Austrian government.

Pertaining to the preparation of neuronal cells from Sprague-Dawley rats at the Mainz facility, all animal procedures were conducted in accordance with German federal law, adhering to the institutional guidelines of the Central Animal Facility of the University of Mainz.

### Brain region-specific tissue preparation

#### Single treatment study

To separate different brain regions of interest, whole brains were sectioned into slices of 750 µm thickness using a vibratome. Therefore, the frozen brains were fixed on a specimen holder with tissue glue, stabilized with an agarose block and sliced while immerged in ice-cold Ringer solution (125 mM NaCl, 2.5 mM KCl, 1.25 mM NaH_2_PO_4,_ 25 mM D-Glucose, 1 mM MgCl_2,_ 2 mM CaCl_2_ 25 mM HEPES, pH 7.4 with NaOH, ultra-pure water). The brains were sliced into coronal sections from rostal to caudal. Each slice was transferred to a small cell culture plate placed on ice and filled with ice-cold Ringer solution supplemented with phosphatase inhibitors (2 mM imidazole, 1 mM NaF, 1 mM Na_4_VO_4_). The brain regions were sampled from these slices by visually dissecting the appropriate coronal and horizontal Bregma parts under a binocular, following the definitions set forth in the literature (Paxinos and Franklin, 2001). The hippocampus (Hc) and the cortex (Ctx) were collected from coronal Bregma -1.06 to -3.08; the collected region termed “cortex” contained the pooled areas of the somatosensory, visual, auditory and ectorhinal cortices and was separated from the regions of the retrosplenial cortex (Rsp) and the entorhinal cortex (Ent), which were sampled independently. The retrosplenial cortex was collected from coronal Bregma -1.06 to -3.08; the entorhinal cortex (encompassing a certain amount of the adjacent piriform cortex, but without the subiculum) was collected from coronal Bregma -2.54 to -5.20. The thalamus (Th) was collected from coronal Bregma -1.06 to -2.70; the regions of the caudate and the putamen (CPu) were dissected from coronal Bregma 1.54 to -0.94. All samples were lysed in sucrose- and SDS-containing buffer (100 mM Tris, 20% sucrose, 0.5% SDS, pH 7.4 with HCl) supplemented with 1x protease and 1x phosphatase inhibitor cocktails (P8340 from Sigma-Aldrich; PhosSTOP from Roche). The sectioned tissues pieces were mechanically homogenized, briefly sonicated and stored at -80°C for further analysis.

#### Repeated treatment study

The frozen brains were carefully cut into two halves at the median sagittal plane. The left hemisphere was used for tissue preparation and was therefore immersed in ice-cold Ringer solution (125 mM NaCl, 2.5 mM KCl, 1.25 mM NaH_2_PO_4,_ 25 mM D-Glucose, 1 mM MgCl_2,_.2 mM CaCl_2,_ 25 mM HEPES, pH 7.4 with NaOH, ultra-pure water). Using a binocular, the areas of the hippocampus, the cortex, the thalamus and the cerebellum were separated according to their visible anatomical boundaries (Paxinos and Franklin, 2001). The tissues were lysed in 100 mM Tris buffer (pH 7.4 with HCl) containing 1x protease and 1x phosphatase inhibitor cocktails (P8340 from Sigma-Aldrich; PhosSTOP from Roche). The pieces of tissue were gently crushed with a pistil and further homogenized by sonication before being stored at -80°C. For the subsequent Western blot analyses, the samples were diluted in sucrose- and SDS-containing buffer (100 mM Tris, 20% sucrose, 0.5% SDS, pH 7.4 with HCl) supplemented with the same protease and phosphatase inhibitor cocktails.

### Immunoblotting

Prior to immunoblotting, the total protein content of all samples was determined by the BCA method using commercial components (Thermo Fisher Scientific). Proteins were separated either by freshly polymerized 10% SDS-polyacrylamide gels or by precast 10-20% Tricine-SDS-polyacrylamide gradient gels (Thermo Fisher Scientific). Gradient gels were exclusively used to separate APP and its cleavage fragments following a published protocol (Rosen et al., 2010), but adopting a different antibody (see below) to detect endogenous mouse and rat APP. For all other purposes, isocratic 10% gels were used. For the SDS-PAGE, samples of 10 µg protein were diluted with 0.25 volumes of 4x sample buffer (200 mM Tris/HCl pH 8.6, 40% glycerol, 20 mM EDTA, 0.08% bromophenol blue, 8% SDS, 20% mercaptoethanol) and boiled for 5 min at 95°C. After loading the gel, electrophoresis was performed at 120 V for 1.5 h. The separated proteins were immediately transferred to a 0.2 µm nitrocellulose membrane via overnight electroblotting at 30 V. On the next day, the membranes were stained (0.2% Ponceau S in 5% acetic acid), photographed, and subsequently blocked with 2% non-fat dry milk (Carl Roth) in TBST (TBS with 0.05% Tween 20) for 60 min. After three washes with TBST, the membranes were incubated with a primary antibody at 4°C overnight.

Antibodies were diluted in TBST supplemented with 0.04% NaN_3_ (except for HRP-conjugated antibodies). The following antibodies were used: anti-Actin clone AC-40 (A-4700 from Sigma-Aldrich, dilution 1:1000); anti-ADAM10 (AB19026, lots #0702052630 and 2776884 from Merck (Darmstadt, Germany), dilution 1:500); anti-APP (ab2072, lots #GR25367-12 and #GR25367-13 from abcam (Cambridge, UK), dilution 1:500); anti-AT8 (MN1020, lots #QE217252 and #SJ2448082 from Thermo Fisher Scientific, dilution 1:200); anti-BACE1 (B0681, lot #076K4780 from Sigma-Aldrich, dilution 1:1000); anti-PS1 (3622, lot #2 from Cell Signaling (Danvers, MA, USA), dilution 1:500); anti-Tau-1 clone PC1C6 (MAB3420, lots #254903 and #2683924 from Merck, dilution 1:500); anti-Tau phospho-S404 (ab92676, lot #GR45351-13 from abcam, dilution 1:1000); anti-Tau phospho-S396 (ab109390, lot #GR189096-9 from abcam, dilution 1:3000); anti-αTub clone DM1A, HRP-conjugated (ab40742, lots #GR174230-1 and GR314710-1 from abcam, 1:3000). After removal of the primary antibody, the membranes were washed three times with TBST and incubated for 2 h at room temperature with an appropriate HRP-conjugated secondary antibody (from Dianova, Hamburg, Germany) diluted 1:10.000 in TBST. Band detection was done by enhanced chemiluminescence. Solution A (100 mM Tris/HCl (pH 8.6), 0.05% luminol) was freshly mixed with solution B (0.11% p-hydroxycoumaric acid) at a ratio of 10:1 before adding 10% of a concentrated H_2_O_2_ stock solution (30% w/v). The membrane was rapidly and consistently coated with the resulting mixture, and the emitted light was recorded with an optical imager. Protein band intensities were quantified using commercial image analysis software (AIDA Image Analyzer from Raytest, Straubenhardt, Germany).

Western blot analyses of cell culture experiments were done on three independent biological replicates (n = 3 cell cultures from three separate litters) for each K+ condition. The analysis of *Single treatment study*-mice was done on five animals out of each group (n = 5) to fit the employed 15-lane gels. The numerically first 5 animals out of the 7-member saline control group were arbitrarily chosen for analysis and retained throughout all Western blot experiments. For the same reason (15-lane gels), only the first 7 mice out of the 8-membered groups of the *Repeated treatment study* were analyzed when conducting Western blot experiments.

### Amyloid β quantification

To determine the amount of Aβ_40_ and Aβ_42_ in vitro and in vivo, commercial ELISA kits were used (Human/Rat β Amyloid (40) ELISA Kit, 294-62501, lots #WDP2982 and #WDL6357, and Human/Rat β Amyloid (42) ELISA Kit High-Sensitive, 292-64501, lots #WDQ1686 and #WDK0714, both from Wako Pure Chemicals, Osaka, Japan). These ELISAs have been shown to be superior to many other antibody combinations (signal-to-noise ratios >15 when testing for endogenous murine amyloid) and have been certified to be suitable for the detection of endogenous levels of Aβ _40_ and Aβ _42_ (Teich et al., 2013).

For the measurement of Aβ secretion from cultivated cortical cells, the experimental medium (Ringer solution) after 4 h treatment with glutamatergic agonists and antagonists was collected, supplemented with 1x protease and 1x phosphatase inhibitor cocktails (P8340 from Sigma-Aldrich; PhosSTOP from Roche) and stirred for 10 min at 37°C. Afterwards, the samples were centrifuged at 20,000 g for 10 min at room temperature. The resulting supernatant was used to perform the ELISAs with 100 µl per well following the manufacturer’s instructions.

The single-treatment in vivo samples that had been lysed in SDS-containing buffer (100 mM Tris, 20% sucrose, 0.5% SDS, pH 7.4 with HCl) were subjected to purification by either of two methods to reduce the amount of free SDS. Method #1 was applied to hippocampal and cortical samples and involved the application of the samples to commercial detergent removal spin columns (87777 from Thermo Fisher Scientific). After adjustment to an equal protein concentration of 2 mg/ml with the original tissue lysis buffer, the samples were centrifuged at 1,500 g for 2 min at room temperature as per the manufacturer’s instructions. The obtained, SDS-depleted samples were then diluted to a final working concentration of 1 mg/ml with the “standard diluent” provided as part of each ELISA Kit. Method #2 was applied to cortical samples (for control purposes) as well as to samples from the thalamus, the entorhinal cortex, the retrosplenial cortex and the caudate/putamen. Here, SDS reduction was achieved by precipitation with KCl under conditions that have been shown to yield high peptide recovery (Zhou et al., 2012). Samples were adjusted to a protein concentration of 2 mg/ml with the original lysis buffer before addition of 200 mM KCl from a 2 M stock in ultra-pure water. After 10 min incubation at room temperature, the KDS precipitate was removed by centrifugation at 15,000 g for 10 min. The obtained supernatant was diluted 1:1 with ultra-pure water to obtain the final working concentration of 1 mg/ml protein. ELISAs were then performed as per the manufacturer’s instructions. Comparison of the two SDS reduction protocols was done by inspecting the ELISA results for the cortex, which was analyzed by both protocols. The obtained values for Aβ _40_ and Aβ _42_ according to both protocols were very similar, such that the results from method #2 were arbitrarily chosen for presentation. In ELISAs with material from the in vivo studies, all n = 7 saline control mice were analyzed.

## Statistical analysis

In vitro experiments were statistically evaluated by parametric Two-way ANOVAs. Pertaining to all these Two-way ANOVAs, factor 1 (indexed as subscript1) relates to excitatory agonist exposure (NMDA or Glu), and factor 2 (indexed as subscript_2_) relates to excitatory antagonist (AP5) exposure. In vivo studies were evaluated by parametric One-way ANOVAs comparing saline-treated control animals with MK-801 treated animals. To identify pairwise significant differences versus the control after Two-way or One-way ANOVAs, Holm-Sidak’s post-hoc test was employed. All significant p values (versus the control at the p = 0.05 level) resulting from these post-hoc tests are provided in the figures. All p values in the main text and the figure legend text represent the corresponding maternal ANOVA significances. The Supplemental information provides a complete list of all maternal ANOVA significances including F values. In cases where the ANOVA normality assumption or equal variance assumption were not met at the p = 0.05 level (normality test failed, n.t.f.; equal variance test failed, e.v.f.), the provided Two-way or One-way ANOVA results are complemented (in the Supplemental information) with the results of One-way ANOVA on Ranks evaluations. All data in this work are shown as mean and standard deviation.

## Supplemental information

### Statistical details

Statistical evaluations were generally done as parametric Two-way ANOVA or parametric One-way ANOVA as indicated. Pertaining to all Two-way ANOVAs, factor 1 (indexed as subscript1) relates to excitatory agonist exposure (NMDA or Glu), and factor 2 (indexed as subscript_2_) relates to excitatory antagonist (AP5) exposure.

In cases where the normality assumption or the equal variance assumption were not met at the p = 0.05 level (normality test failed, n.t.f.; equal variance test failed, e.v.f.), the provided Two-way or One-way ANOVA results are complemented with the results of One-way ANOVA on Ranks evaluations, which are given in brackets below each of the retested parametric ANOVAs.

#### S1: Statistical details Figure 1

Evaluations were done by Two-way ANOVA.

**(b)**

**low K+:**

anti-AT8: F_1_ = 1.580, d.f. = 2, p_1_ = 0.246; F_2_ = 22.825, d.f. = 1, p_2_ < 0.001; n = 3;

anti-S404: F_1_ = 1.608, d.f. = 2, p_1_ = 0.241; F_2_ = 12.531, d.f. = 1, p_2_ = 0.004; n = 3;

anti-S396: F_1_ = 0.489, d.f. = 2, p_1_ = 0.625; F_2_ = 7.608, d.f. = 1, p_2_ = 0.017; n = 3; n.t.f.;

[anti-S396: H = 7.111, d.f. = 5, p = 0.212; n = 3].

**high K+:**

anti-AT8: F_1_ = 0.797, d.f. = 2, p_1_ = 0.473; F_2_ = 6.660, d.f. = 1, p_2_ = 0.024; n = 3;

anti-S404: F_1_ = 1.245, d.f. = 2, p_1_ = 0.322; F_2_ = 40.611, d.f. = 1, p_2_ < 0.001; n = 3;

anti-S396: F_1_ = 0.316, d.f. = 2, p_1_ = 0.735; F_2_ = 7.731, d.f. = 1, p_2_ = 0.017; n = 3.

**(c)**

**low/high K+ combined:**

anti-AT8: F_1_ = 2.263, d.f. = 2, p_1_ = 0.122; F_2_ = 26.695, d.f. = 1, p_2_ < 0.001; n = 6;

anti-S404: F_1_ = 2.034, d.f. = 2, p_1_ = 0.148; F_2_ = 43.670, d.f. = 1, p_2_ < 0.001; n = 6; e.v.f.;[anti-S404: H = 22.100, d.f. = 5, p < 0.001; n = 6];

anti-S396: F_1_ = 0.948, d.f. = 2, p_1_ = 0.399; F_2_ = 18.806, d.f. = 1, p_2_ < 0.001; n = 6; n.t.f.;[anti-S396: H = 17.523, d.f. = 5, p = 0.004; n = 6].

**(e)**

**low K+:**

F_1_ = 2.506, d.f. = 2, p_1_ = 0.110; F_2_ = 4.963, d.f. = 1, p_2_ = 0.039; n = 4.

**high K+:**

F_1_ = 2.999, d.f. = 2, p_1_ = 0.075; F_2_ = 0.201, d.f. = 1, p_2_ = 0.659; n = 4.

**(f)**

**low K+:**

F_1_ = 3.784, d.f. = 2, p_1_ = 0.042; F_2_ = 3.811, d.f. = 1, p_2_ = 0.067; n = 4; n.t.f.;[H = 8.306, d.f. = 5, p = 0.140; n = 4].

**high K+:**

F_1_ = 6.184, d.f. = 2, p_1_ = 0.009; F_2_ = 3.939, d.f. = 1, p_2_ = 0.063; n = 4.

#### S2: Statistical details Figure 2

Evaluations were done by Two-way ANOVA.

**(b)**

**low K+:**

F_1_ = 1.787, d.f. = 2, p_1_ = 0.209; F_2_ = 31.863, d.f. = 1, p_2_ < 0.001; n = 3.

**high K+:**

F_1_ = 0.012, d.f. = 2, p_1_ = 0.988; F_2_ = 6.438, d.f. = 1, p_2_ < 0.001; n = 3.

**(c)**

**low/high K+ combined:**

F1 = 1.367, d.f. = 2, p_1_ = 0.270; F_2_ = 89.790, d.f. = 1, p_2_ < 0.001; n = 6; e.v.f.; [H = 28.581, d.f. = 5, p < 0.001; n = 6].

**(d)**

**low/high K+ combined:**

**Aβ40:**

F_1_ = 2.493, d.f. = 2, p_1_ = 0.100; F_2_ = 10.392, d.f. = 1, p_2_ = 0.003; n = 6; n.t.f.;[H = 13.533, d.f. = 5, p = 0.019; n = 6].

**Aβ42:**

F_1_ = 0.245, d.f. = 2, p_1_ = 0.784; F_2_ = 0.009, d.f. = 1, p_2_ = 0.925; n = 6; n.t.f.;[H = 5.230, d.f. = 5, p = 0.388; n = 6].

#### S3: Statistical details Figure 3

Evaluations were done by One-way ANOVA.

**(b)**

**Hippocampus:**

anti-AT8: F = 2.120, d.f. = 2, p = 0.163; n = 5;

anti-S404: F = 5.327, d.f. = 2, p = 0.022; n = 5; n.t.f.;[anti-S404: H = 7.220, d.f. = 2, p = 0.027; n = 5];

anti-S396: F = 0.582, d.f. = 2, p = 0.574; n = 5.

**(d)**

**Cortex:**

anti-AT8: F = 9.998, d.f. = 2, p = 0.003; n = 5;

anti-S404: F = 5.970, d.f. = 2, p = 0.016; n = 5;

anti-S396: F = 2.388, d.f. = 2, p = 0.134; n = 5.

**Thalamus:**

anti-AT8: F = 0.068, d.f. = 2, p = 0.935; n = 5;

anti-S404: F = 1.137, d.f. = 2, p = 0.353; n = 5;

anti-S396: F = 2.470, d.f. = 2, p = 0.126; n = 5.

#### S4: Statistical details Figure 4

Evaluations were done by One-way ANOVA.

**(d)**

**Hippocampus:**

F = 11.945, d.f. = 2, p = 0.001; n = 5.

**Cortex:**

F = 7.635, d.f. = 2, p = 0.007; n = 5.

**Thalamus:**

F = 4.207, d.f. = 2, p = 0.041; n = 5.

**(e)**

**Hippocampus:**

F = 4.588, d.f. = 2, p = 0.029; n = 7 for Ctrl, n = 5 for 4 h and 24 h.

**Cortex:**

F = 2.312, d.f. = 2, p = 0.136; n = 7 for Ctrl, n = 5 for 4 h and 24 h.

**Thalamus:**

F = 2.585, d.f. = 2, p = 0.111; n = 7 for Ctrl, n = 5 for 4 h and 24 h.

**Entorhinal cortex:**

F = 0.536, d.f. = 2, p = 0.597; n = 7 for Ctrl, n = 5 for 4 h and 24 h; n.t.f.;

[H = 1.294, d.f. = 2, p = 0.524; n = 7 for Ctrl, n = 5 for 4 h and 24 h].

**Retrosplenial cortex:**

F = 5.690, d.f. = 2, p = 0.016; n = 7 for Ctrl, n = 5 for 4 h and 24 h; n.t.f.;

[H = 7.228, d.f. = 2, p = 0.027; n = 7 for Ctrl, n = 5 for 4 h and 24 h].

**Caudate/putamen:**

F = 2.440, d.f. = 1, p = 0.149; n = 7 for Ctrl, n = 5 for 4 h; n.t.f.;

[H = 2.656, d.f. = 1, p = 0.106; n = 7 for Ctrl, n = 5 for 4 h].

**(f)**

**Hippocampus:**

F = 6.509, d.f. = 2, p = 0.010; n = 7 for Ctrl, n = 5 for 4 h and 24 h.

**Cortex:**

F = 4.355, d.f. = 2, p = 0.034; n = 7 for Ctrl, n = 5 for 4 h and 24 h.

**Thalamus:**

F = 13.313, d.f. = 2, p < 0.001; n = 7 for Ctrl, n = 5 for 4 h and 24 h.

**Entorhinal cortex:**

F = 1.460, d.f. = 2, p = 0.266; n = 7 for Ctrl, n = 5 for 4 h and 24 h.

**Retrosplenial cortex:**

F = 0.034, d.f. = 2, p = 0.966; n = 7 for Ctrl, n = 5 for 4 h and 24 h; n.t.f.;[H = 0.070, d.f. = 2, p = 0.966; n = 7 for Ctrl, n = 5 for 4 h and 24 h].

**Caudate/putamen:**

F = 2.645, d.f. = 1, p = 0.135; n = 7 for Ctrl, n = 5 for 4 h; n.t.f.;[H = 1.905, d.f. = 1, p = 0.202; n = 7 for Ctrl, n = 5 for 4 h].

**(g)**

**Hippocampus:**

F = 5.754, d.f. = 2, p = 0.015; n = 7 for Ctrl, n = 5 for 4 h and 24 h.

**Cortex:**

F = 1.168, d.f. = 2, p = 0.340; n = 7 for Ctrl, n = 5 for 4 h and 24 h.

**Thalamus:**

F = 5.924, d.f. = 2, p = 0.014; n = 7 for Ctrl, n = 5 for 4 h and 24 h.

**Entorhinal cortex:**

F = 0.426, d.f. = 2, p = 0.661; n = 7 for Ctrl, n = 5 for 4 h and 24 h; n.t.f.;

[H = 1.425, d.f. = 2, p = 0.490; n = 7 for Ctrl, n = 5 for 4 h and 24 h].

**Retrosplenial cortex:**

F = 5.299, d.f. = 2, p = 0.019; n = 7 for Ctrl, n = 5 for 4 h and 24 h.

**Caudate/putamen:**

F = 5.335, d.f. = 1, p = 0.044; n = 7 for Ctrl, n = 5 for 4 h; n.t.f.;[H = 4.121, d.f. = 1, p = 0.048; n = 7 for Ctrl, n = 5 for 4 h].

**(h)**

**Hippocampus:**

anti-ADAM10: F = 2.621, d.f. = 2, p = 0.114; n = 5;

anti-BACE1: F = 0.334, d.f. = 2, p = 0.722; n = 5;

anti-PS1: F = 5.233, d.f. = 2, p = 0.023; n = 5.

**Cortex:**

anti-ADAM10: F = 5.027, d.f. = 2, p = 0.026; n = 5;

anti-BACE1: F = 2.056, d.f. = 2, p = 0.171; n = 5;

anti-PS1: F = 6.826, d.f. = 2, p = 0.010; n = 5.

**Thalamus:**

anti-ADAM10: F = 4.132, d.f. = 2, p = 0.043; n = 5;

anti-BACE1: F = 1.735, d.f. = 2, p = 0.218; n = 5; anti-PS1: F = 0.412, d.f. = 2, p = 0.671; n = 5; n.t.f.;

[anti-PS1: H = 1.460, d.f. = 2, p = 0.482; n = 5].

#### S5: Statistical details Figure 5

Evaluations were done by One-way ANOVA.

**(b)**

**Hippocampus:**

anti-AT8: F = 1.011, d.f. = 1, p = 0.335; n = 7;

anti-S404: F = 0.371, d.f. = 1, p = 0.554; n = 7;

anti-S396: F = 0.000, d.f. = 1, p = 0.996; n = 7; e.v.f.;

[anti-396: H = 0.004, d.f. = 1, p = 1.000; n = 7].

**(d)**

**Cortex:**

anti-AT8: F = 5.395, d.f. = 1, p = 0.039; n = 7;

anti-S404: F = 9.143, d.f. = 1, p = 0.011; n = 7;

anti-S396: F = 0.054, d.f. = 1, p = 0.819; n = 7.

**(e)**

**Hippocampus:**

F = 0.898, d.f. = 1, p = 0.362; n = 7; n.t.f.;[H = 0.102, d.f. = 1, p = 0.805; n = 7].

**Cortex:**

F = 5.319, d.f. = 1, p = 0.039; n = 7.

#### S6: Statistical details Figure 6

Evaluations were done by One-way ANOVA.

**(b)**

**Hippocampus:**

anti-AT8: F = 13.055, d.f. = 1, p = 0.004; n = 7; e.v.f.;[anti-AT8: H = 6.208, d.f. = 1, p = 0.011; n = 7];

anti-S404: F = 12.466, d.f. = 1, p = 0.004; n = 7;

anti-S396: F = 1.827, d.f. = 1, p = 0.201; n = 7.

**(d)**

**Cortex:**

anti-AT8: F = 1.704, d.f. = 1, p = 0.216; n = 7; e.v.f.;[anti-AT8: H = 0.918, d.f. = 1, p = 0.383; n = 7];

anti-S404: F = 16.652, d.f. = 1, p = 0.002; n = 7;

anti-S396: F = 1.445, d.f. = 1, p = 0.252; n = 7.

**(e)**

**Hippocampus:**

F = 2.066, d.f. = 1, p = 0.176; n = 7.

**Cortex:**

F = 0.704, d.f. = 1, p = 0.418; n = 7.

**(f)**

**Hippocampus:**

anti-ADAM10: F = 0.004, d.f. = 1, p = 0.949; n = 7;

anti-BACE1: F = 0.654, d.f. = 1, p = 0.434; n = 7;

anti-PS1: F = 6.026, d.f. = 1, p = 0.030; n = 7.

**Cortex:**

anti-ADAM10: F = 0.012, d.f. = 1, p = 0.916; n = 7;

anti-BACE1: F = 0.374, d.f. = 1, p = 0.552; n = 7;

anti-PS1: F = 7.437, d.f. = 1, p = 0.018; n = 7.

#### S7: Statistical details Figure 7

Evaluations were done by One-way ANOVA.

**(b)**

**Cortex:**

anti-AT8: F = 16.543, d.f. = 1, p = 0.002; n = 7;

anti-S404: F = 12.572, d.f. = 1, p = 0.004; n = 7; n.t.f.;[anti-S404: H = 8.265, d.f. = 1, p = 0.002; n = 7];

anti-S396: F = 9.747, d.f. = 1, p = 0.009; n = 7; e.v.f.;[anti-S396: H = 6.208, d.f. = 1, p = 0.011; n = 7].

